# Unravelling differential Hes1 dynamics during axis elongation of mouse embryos through single-cell tracking

**DOI:** 10.1101/2024.03.30.586691

**Authors:** Yasmine el Azhar, Pascal Schulthess, Marek J. van Oostrom, Wilke H.M. Meijer, Wouter M. Thomas, Marianne Bauer, Katharina F. Sonnen

## Abstract

The intricate dynamics of Hes expression across diverse cell types in the developing vertebrate embryonic tail have remained elusive. To address this, we developed an endogenously tagged Hes1-Achilles mouse line, enabling precise quantification of dynamics at the single-cell resolution across various tissues. Our findings reveal striking disparities in Hes1 dynamics between presomitic mesoderm (PSM) and preneural tube (pre-NT) cells. While pre-NT cells display variable, low-amplitude oscillations, PSM cells exhibit synchronized, high-amplitude oscillations. Upon the induction of differentiation, the oscillation amplitude increases in pre-NT cells. Additionally, our study of Notch inhibition on Hes1 oscillations unveiled distinct responses in PSM and pre-NT cells, corresponding to differential Notch ligand expression dynamics. These findings suggest the involvement of separate mechanisms driving Hes1 oscillations. Thus, Hes1 demonstrates dynamic behaviour across adjacent tissues of the embryonic tail, yet the varying oscillation parameters give rise to differences in the information that can be conveyed by these dynamics.

## Introduction

During embryonic development, tightly regulated cell fate decisions rely on intercellular communication. Recent insights indicate that cells encode information not only through the presence but also the dynamics of signalling pathways^1^. Oscillations in signalling pathways and target genes are widespread across tissues and developmental stages. Prominent among these are members of the hairy and enhancer of split (Hes) family of transcriptional repressors. Hes proteins, basic helix-loop-helix (bHLH) transcription factors, modulate gene expression by interacting with DNA as dimers^2^. Several of the seven Hes family members in mammalian cells are dynamically expressed in embryonic development^2-7^.

During vertebrate development, neuromesodermal progenitors (NMPs) in the tailbud (TB) give rise to both the mesodermal and neural lineages, namely the presomitic mesoderm (PSM) and the neural tube (NT), respectively^8-10^. Hes dynamics play a critical role in the differentiation of both PSM and NT progenitors^3,11-13^. The periodic segmentation of the PSM into somites, known as somitogenesis, is driven by oscillatory signalling pathways, including FGF (fibroblast growth factor), Wnt (wingless and Int-1) and Notch signalling^4,14-18^. Whereas the exact composition of the segmentation clock differs per species, Notch signalling oscillations are found in all studied vertebrates, and several Notch target genes display oscillatory expression in PSM cells^4,17,19-22^, including several members of the Hes gene family^4,13,23,24^.

NMPs transitioning from the TB to the NT pass through the preneural tube (pre-NT), surrounded by oscillating PSM^25^. FGF signalling prevents premature differentiation in the pre-NT, while Notch signalling supports cell proliferation^26,27^. In the anterior NT, progenitor cells continue to proliferate and differentiate into future spinal cord neurons. Whether NT progenitor cells proliferate or differentiate has been linked to oscillations in Hes1 and Hes5, reminiscent of their function in somitogenesis^11,12^. Hes genes oscillate in progenitor cells to maintain a proliferative state^11,12,28-30^, while sustained expression has been linked to quiescence^31,32^. Despite research into Hes dynamics in the anterior NT, investigations into Hes dynamics in NMPs and pre-NT are lacking.

In this study, we delineate the dynamics of Hes gene expression in mouse development and elucidate the signalling pathways orchestrating its expression during the differentiation of NMPs along neural and mesodermal trajectories. Leveraging a newly developed homozygously viable Hes1 reporter mouse line, generated through the endogenous tagging of the gene locus with a rapidly maturing yellow fluorescent protein, we thoroughly quantify Hes1 dynamics in the PSM, TB and pre-NT regions of the embryonic tail. Through single-cell tracking, we discern distinct Hes1 dynamics between the PSM and pre-NT. Further investigation into Hes1 dynamics revealed disparate responses of PSM and pre-NT cells to signalling pathway perturbations. This also correlated with changes in proliferation. Thus, our new endogenous reporter mouse line, combined with recently available single-cell tracking software, allowed us to uncover previously unquantified dynamics in single cells of the mouse embryonic tail.

## Results

### Generation of a Hes1-Achilles mouse line to study signalling dynamics in the posterior embryonic tail

To investigate Hes dynamics in the developing mouse embryo, we first clarified which Hes genes were expressed in the PSM, TB and in particular the pre-NT. The latter being defined as the tube-like Pax6-negative region emanating from the TB, which lies next to the PSM (Fig. 1A, B)^26,27^. To this end, we performed *in situ* hybridization (ISH) and hybridization chain reactions (HCR)^33^. Expression of Hes7 and Lfng (lunatic fringe), key components of the segmentation clock^24,34-37^, was restricted to the PSM and TB or PSM, respectively (Fig. S1A, B, D). In contrast, both Hes1 and Hes5 were present in the TB, the PSM and the pre-NT with highest expression levels in the pre-NT and TB (Fig. 1C, S1C, D) (also see MAMEP database, http://mamep.molgen.mpg.de). Thus, Hes1 and Hes5 are not limited to PSM, but are also expressed in the TB and pre-NT of the embryonic tail.

**Figure 1.**
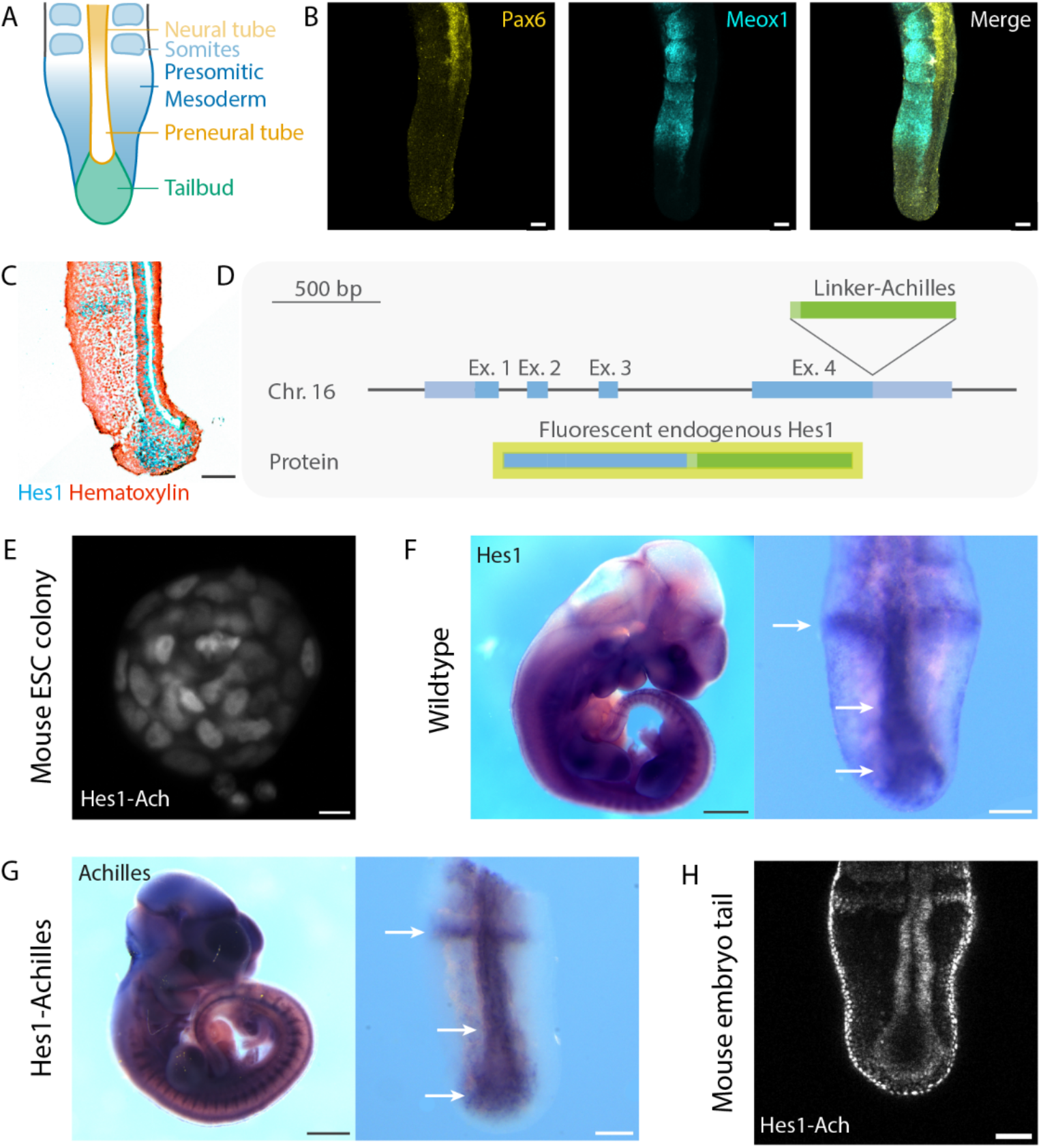
Generation of a Hes1-Achilles mouse line to study Hes dynamics in the posterior embryonic tail. A. Schematic representation of the posterior mouse embryonic tail. Tailbud (green) differentiate into PSM (blue), which gives rise to somites (light blue), and the pre-NT (yellow), which differentiates into the NT (light yellow). **B** Representative image of a E10.5 embryonic tail stained by HCR for the mesodermal marker Meox1 and neural marker Pax6. Scale bar is 100 µm. **C** 5 µm section of an ISH of an E10.5 embryonic tail for Hes1 (as shown in Fig. S1F). Scale bar is 100 µm. **D** Schematic representation of the tagging strategy for the Hes1-Achilles mouse line. Hes1 was endogenously tagged with linker-Achilles. **E** Representative image of a mouse ESC colony expressing the Hes1-Achilles construct. Note the variable expression in the different cell types. Scale bar is 10 µm. **F, G** Representative images of ISH of an E10.5 embryo for Hes1 (**F**) (as shown in Fig. S1F) and Achilles (**G**). Right panel shows magnification of the posterior embryonic tail. Arrows highlight the expression pattern in the posterior embryonic tail. Scale bar is 100 µm. **H** Representative image of an E10.5 embryonic tail expressing the Hes1-Achilles reporter. Scale bar is 100 µm.

For further analysis into the dynamics of Hes genes, we then focused on Hes1, which shows a more localized expression to the neural lineage and PSM of the developing embryo than the ubiquitously expressed Hes5 (MAMEP database). To be able to quantify expression levels over time, we generated a new Hes1 mouse line (Fig. 1D-H). Building on previous approaches^31,38-41^, we endogenously tagged Hes1 with Achilles, a fast-maturing fluorescent protein^39^ (Fig. 1D, S2). We placed Achilles at the C-terminal side of Hes1 as this lies opposite to the DNA-binding bHLH domain of Hes1^2^. In addition, we separated Hes1 and Achilles by a flexible linker, as we reasoned that this should allow interaction of Hes1 via its C-terminal WRPW motif (four amino acid protein-protein interaction domain) with the Groucho/TLE family of transcriptional repressors^42,43^. Structure of the fusion protein was assessed for accessibility of the WRPW motif and bHLH domain by Alphafold^44^ (data not shown). Homozygous mice were born at Mendelian ratio (offspring of 8 heterozygous breedings: 17 wildtype, 34 heterozygous, 16 homozygous), viable and fertile. When comparing mRNA staining of Hes1 in wildtype embryos and Achilles in Hes1-Achilles embryos, the expression patterns corresponded (Fig. 1F, G). At protein level, Hes1-Achilles expression indicated the expected pattern with high expression levels in the pre-NT and stripes in forming somites (Fig. 1C, H). Thus, we successfully generated a homozygously viable, fertile and endogenous Hes1-Achilles reporter mouse line.

### Hes1 expression is dependent on Notch signalling in the posterior embryonic tail

We next addressed whether Hes1 expression was solely dependent on Notch signalling or also on other pathways, as suggested previously for Hes7^40^. To be able to test this, we had to culture E10.5 embryonic tails *ex vivo* to quantify Hes1-Achilles dynamics upon pathway perturbation. However, standard embryo culture protocols^45,46^ needed to be optimized for survival of the NT. We did this by testing different media to improve pre-NT and NT survival (Fig. 2A and data not shown). In standard embryo culture medium, cleaved caspase-3 expression levels were significantly higher than in uncultured embryos at embryonic day E11.5, especially in the pre-NT and NT (Fig. 2A, B). The tube was often filled with dead cells or disintegrated entirely. In contrast, in embryonic tails cultured in neurobasal medium (see further details in the Methods), the number of apoptotic cells was not increased compared to uncultured tails. Furthermore, the NT maintained its morphology and proliferative state (Fig. 2A, C). This suggests that the pre-NT is viable in the neurobasal medium. Despite this, somites were less well organized in neurobasal medium than in control samples (Fig. 2A). To find out whether culture conditions influenced the well-studied dynamics of the segmentation clock, we compared embryonic tails expressing LuVeLu^47^ cultured in either embryo culture medium or neurobasal medium (Fig. S3A-D). Wave and oscillation dynamics as well as the regression of the oscillating field, a sign of differentiation, were similar between embryonic tails cultured in the different media (Fig. S3A, B). Accordingly, period and amplitude remained unchanged (Fig S3C, D). Thus, dynamics of the segmentation clock were maintained, and the optimized culture conditions of embryonic tails now allow the investigation of signalling dynamics in PSM, TB and pre-NT *ex vivo*.

**Figure 2.**
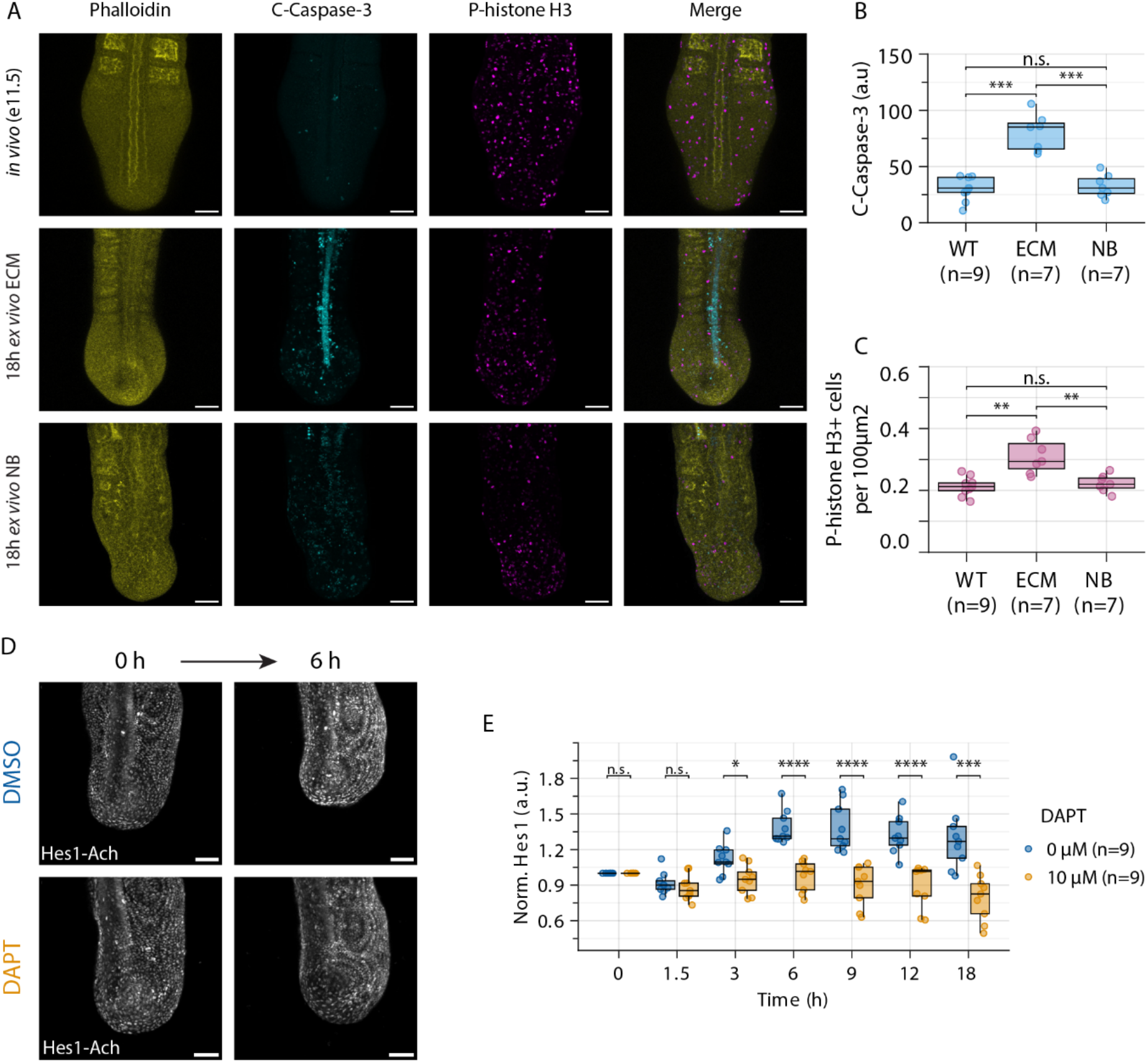
Inhibition of Notch signalling results in a decrease in Hes1-Achilles levels in *ex vivo* cultured E10.5 embryonic tails. A-C. Comparison of different culture media to optimize for the survival of the pre-NT *ex vivo*. Embryonic tails were cultured *ex vivo* for 18 h in embryo culture medium (ECM) or neurobasal (NB) medium and compared to uncultured (*in vivo*) E11.5 embryos. Immunostaining against cleaved-caspase 3 (C-Caspase 3, apoptosis) phospho-histone H3 (P-histone H3, mitosis) and counterstaining with Phalloidin was performed. In **A** representative images are shown. In **B** the intensity of cleaved-caspase 3 signal in the embryonic tail and in **C** the number of mitotic cells in the pre-NT were quantified. **D, E** E10.5 embryonic tails were cultured *ex vivo* and incubated with DMSO control or the gamma-secretase inhibitor DAPT. Hes1-Achilles intensity (normalised to the first timepoint) was measured over time. In **D** representative images and in **E** the quantification of Hes1-Achilles intensity in the embryonic tail are shown. Scale bar is 100 µm. Dots in boxplots represent individual data points. * is p<0.05, ** is p<0.01, *** is p<0.001, **** is p<0.0001.

Using these culture conditions, we initially tested how pathway perturbation affected Hes1-Achilles expression levels. Notch inhibition, using the gamma-secretase inhibitor DAPT, led to a drop in Hes1-Achilles expression levels within 3 h (Fig. 2D, E). The dynamics and signal loss were comparable to the effect of Notch inhibition on the reporter Achilles-Hes7^39^ (Fig. S3F, G). Besides Notch signalling, we also focused on Wnt (Fig. S3H-J) and FGF (Fig. S3K-M) signalling, as gradients of these morphogens are present in the embryonic tail including the pre-NT (Fig. S3E). Accordingly, we found Dusp4 and Axin2, downstream targets of FGF and Wnt signalling, respectively, to be highly expressed in the pre-NT (Fig. S3H, K). We therefore perturbed embryonic tails with small molecules inhibiting these pathways: Wnt signalling was inhibited using the porcupine inhibitor IWP2 and FGF signalling was inhibited using SU5402, a small molecule inhibitor against the FGF receptor. Inhibiting either did not lead to a direct (IWP2) or consistent (SU5402) drop in signal, suggesting that these pathways do not have a direct effect on Hes1 expression levels. Thus, Hes1 expression in the posterior embryonic tail is driven by Notch signalling.

### Population-wide Hes1 dynamics differ between TB, pre-NT and PSM

Subsequently, we carefully quantified Hes1-Achilles dynamics in the different regions of the embryonic tail (TB, PSM and pre-NT) (Fig. 3A-G, movie 1). We quantified dynamics in the TB region (Fig. 3D), which includes the NMPs but presumably also some posterior PSM cells. Every region significantly differed in expression level, in particular the pre-NT region showed higher Hes1-Achilles expression levels compared to the PSM region (Fig. 3H). In both the PSM and TB region, we detected oscillations over time (Fig. 3C, E). In contrast, Hes1-Achilles expression in the pre-NT region was dynamic, albeit noisier and showed less prominent oscillations over time compared to the other regions (Fig. 3G). When analysing the period using wavelet transform^48,49^, there was a slight increase in period in the TB region compared to PSM region (Fig. 3I). A similar trend towards an elevated period was observed for the pre-NT region. Furthermore, we found that the amplitude of the PSM region was significantly higher than in the pre-NT and TB regions (Fig. 3J). Finally, whereas kymographs revealed travelling waves of Hes1-Achilles expression levels in the PSM region (see * in kymograph), no clear population-wide pattern was detected in the pre-NT region (Fig. 3K). Overall, the data suggests that Hes1 is dynamic in the different regions of the mouse embryonic tail. However, Hes1 expression levels in the pre-NT are noisier and less oscillatory compared to the PSM.

**Figure 3.**
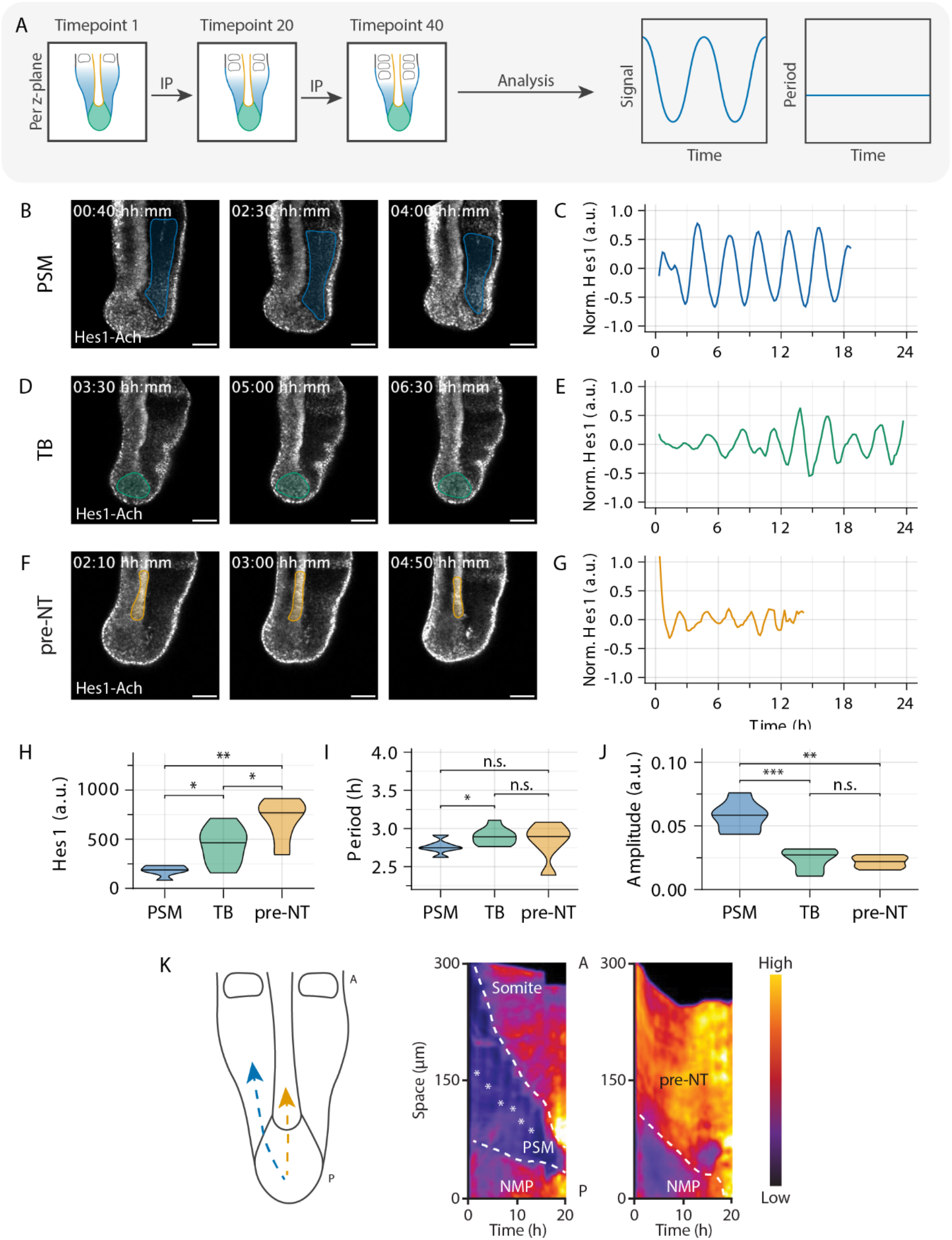
Pre-NT and tailbud region in *ex vivo* cultured E10.5 embryonic tails show low amplitude oscillations at population level. A. Embryonic tails were cultured *ex vivo* and Hes1-Achilles intensity measured by fluorescence real-time imaging (corresponding to correctly positioned control samples from Fig. 2, S3). Regions of interest were selected manually and interpolated (IP) between timepoints. Hes1-Achilles dynamics were then quantified by wavelet transform. **B, C** Quantification of Hes1-Achilles dynamics in the PSM region. Representative snapshots are shown in **B** and the corresponding timeseries data (mean-normalised) of single PSM region (as indicated in **B**) is shown in **C**. **D, E** Quantification of Hes1-Achilles dynamics in the TB region. Representative snapshots are shown in **D** and the corresponding timeseries data (mean-normalised) of single TB region (as indicated in **D**) is shown in in **E**. **F, G** Quantification of Hes1-Achilles dynamics in the pre-NT region. Representative snapshots are shown in **F** and the corresponding timeseries data (mean-normalised) of single pre-NT region (as indicated in **F**) is shown in in **G**. **H** Quantification of absolute Hes1-Achilles intensity in the different regions (PSM: n=6 tails; TB: n=8 tails; pre-NT: n=6 tails). **I, J** Quantification of the period (**I**) and amplitude (**J**) by wavelet transform. **K** Left panel: kymographs of Hes1-Achilles dynamics were generated along the arrows indicated (PSM in blue and pre-NT in yellow) from the posterior (P) to the anterior (A) region of the embryonic tail. Right panel: Representative kymographs for PSM and pre-NT are shown. Hes1-Achilles wave is indicated by asterisk. Scale bar is 100 µm. * is p<0.05, ** is p<0.01, *** is p<0.001, **** is p<0.0001.

### Hes1 is oscillatory in single cells of the pre-NT

The fact that we did not find clear population-wide Hes1 oscillations or travelling waves in the pre-NT region could have different reasons: (1) There are no oscillations. (2) Single cells oscillate but are not synchronized with each other. To test this, we performed a 2D spread-out experiment^46^ combined with mosaic labelling of nuclei that allowed us to track cells individually and quantify Hes1-Achilles expression levels in single cells (Fig. 4A-E, movie 2). When posterior embryonic tail tips are cultured on fibronectin-coated dishes, the tissue spreads and cells grow in a quasi-monolayer, which facilitates single-cell tracking^46^. Immunostaining and HCR for neural and mesodermal markers in 2D cultures confirmed presence and survival of PSM and neural cell types, when culturing in neurobasal medium (Fig. S4A, B). As was observed in the corresponding regions (Fig. 1,3), pre-NT cells showed higher Hes1-Achilles expression levels than PSM cells, the latter of which slightly increased in intensity upon differentiation to somites (Fig. 4F). Interestingly, oscillatory dynamics in Hes1-Achilles were found in both PSM and pre-NT cells (Fig. 4B-E). Wavelet analysis indicated similar wavelet power distributions for PSM and pre-NT cells (data not shown). Likewise, the periods and amplitudes were similar, even though the amplitudes in pre-NT cells were slightly reduced (Fig. 4G, H). The discrepancy between amplitude at population (Fig. 3) and single-cell level (Fig. 4H) supports the hypothesis that Hes1 dynamics are not synchronized between neighbouring pre-NT cells.

**Figure 4.**
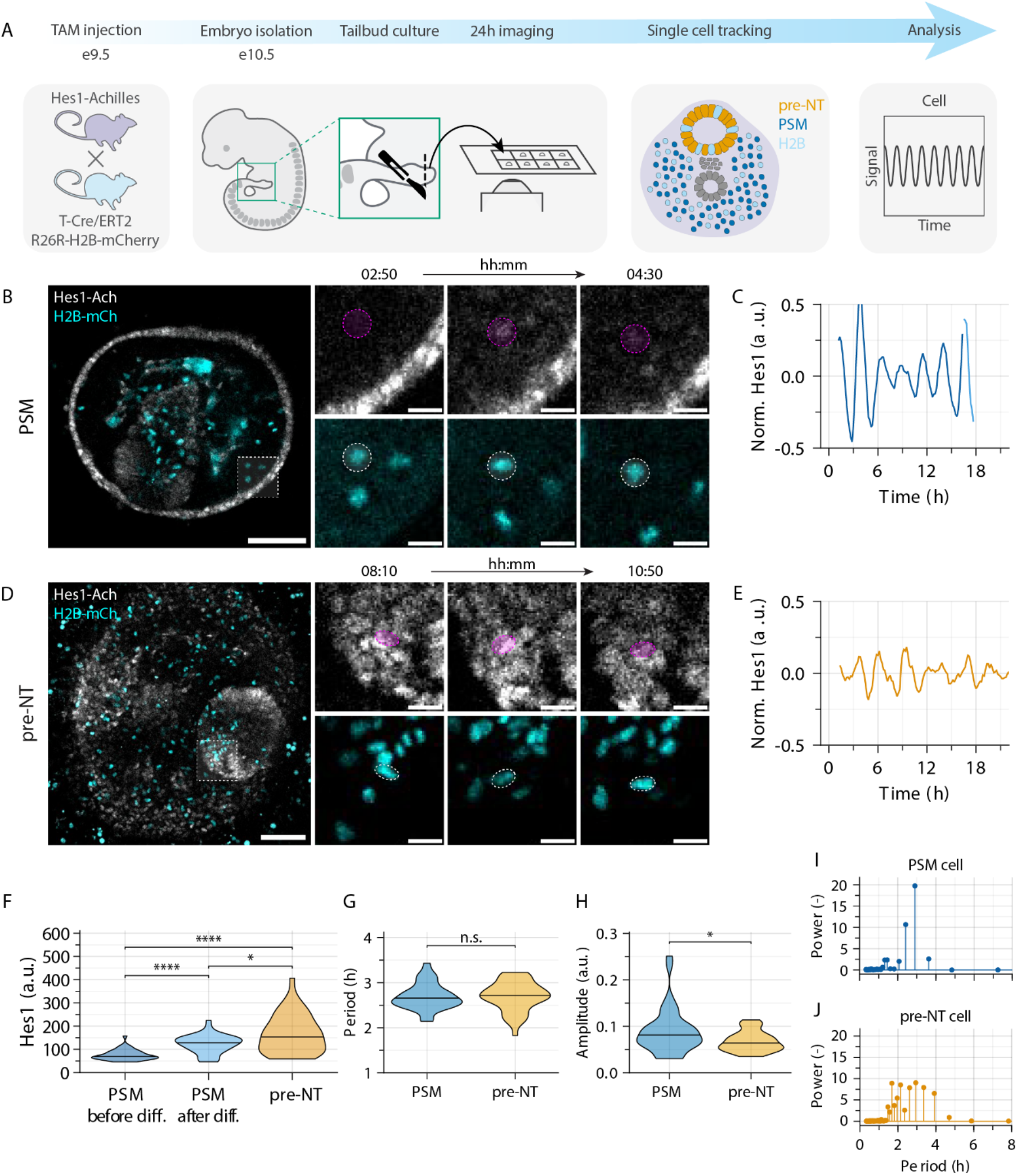
Both PSM and pre-NT cells show Hes1 oscillations at single-cell level. A. Sparse labelling with H2B-mCherry was induced by injection of 1 mg Tamoxifen (TAM). Embryonic tail tips were cultured *ex vivo* on fibronectin-coated dishes. Hes1-Achilles and H2B-mCherry intensity were measured by fluorescence real-time imaging. Single cells were tracked semi-automatically to quantify Hes1-Achilles dynamics. **B, C** Quantification of Hes1-Achilles dynamics in PSM cells. Representative snapshots are shown in **B**. Representative timeseries data (mean-normalised) of a single PSM cell (blue) is shown in **C**. Differentiated PSM cell is indicated by light blue colour. **D, E** Quantification of Hes1-Achilles dynamics in pre-NT cells. Representative snapshots are shown in **D**. Representative timeseries data (mean-normalised) of single pre-NT cell is shown in **E**. **F** Quantification of absolute Hes1-Achilles intensity in the different regions (diff. = differentiation) of the embryonic tail. **G, H** Quantification of the period (**G**) and amplitude (**H**) by wavelet transform. **I, J** Representative Fourier spectrum of a single PSM or pre-NT cell are shown. Note that there is one dominant peak at 2.5 h for PSM, but not one dominant peak for pre-NT. Scale bar is 100 µm. * is p<0.05, ** is p<0.01, *** is p<0.001, **** is p<0.0001. PSM cells: n=39; pre-NT cells: n=40.

Using Fourier transform to analyse the dynamics, similar periods were obtained (Fig. 4I, J, S4C-E). Comparison of Fourier spectra of individual cells indicated that Hes1-Achilles dynamics were noisier in pre-NT than PSM cells, as PSM cells mostly showed one main peak at 2.5 h and pre-NT cells multiple peaks around 2-4 h (Fig. 4I, J, Raw data). As a measure of variability, we determined the full width at half maximum (FWHM) for the period quantification (Fig. S4E), which was 0.5 for the PSM and 1.6 for the pre-NT. The coefficient of variation was 0.1 and 0.2 for PSM and pre-NT cells, respectively. Thus, like PSM cells, single cells in the pre-NT show Hes1 oscillations. However, oscillations are more variable and not synchronized between neighbouring cells.

### Hes1 shows oscillations in NMPs that are driven by Notch signalling

After the discovery of Hes1 dynamics in both PSM and pre-NT cells, we next sought to quantify Hes1 dynamics in NMPs, from which both PSM and pre-NT arise. The 2D spread-out experiments (Fig. 4) did not allow us to unequivocally identify NMPs. Therefore, we analysed Hes1-Achilles dynamics in NMPs by *in vitro* differentiation. To this end, we applied a previously published NMP differentiation protocol (Fig. 5A)^50^ to generate NMPs from embryonic stem cells (ESCs) derived from Hes1-Achilles mice (Fig. 1E). To identify embryonic and neural fate, while quantifying Hes1-Achilles expression levels over time, we added a Sox2-H2B-iRFP reporter (“Sox2-iRFP”) into Hes1-Achilles cells by endogenously tagging Sox2 using CRISPaint^51^. Based on previous literature and immunostainings, we defined the NMP window between 56 and 72 hours (Fig. 5A, S5A)^50,52^. When analysing Hes1-Achilles dynamics at population level, oscillations resembled in appearance and amplitude that of the TB region of the *ex vivo* embryonic tail, albeit with a reduced period (Fig. 5B-E, 3D, E). To find out if single cells show Hes1 oscillations, we next tracked single NMP cells and quantified Hes1-Achilles dynamics (Fig. 5F-I). Hes1-Achilles expression was dynamic in NMPs with a period of around 2.5 h (Fig. 5H). The oscillations were variable and had a low amplitude (Fig. 5G, I). When perturbing Notch signalling using the inhibitor DAPT, we detected a dose-dependent decrease of Hes1-Achilles expression levels (Fig. S5B-E). This inhibition did not show an apparent effect on the period or amplitude of Hes1-Achilles dynamics. Furthermore, when we differentiated NMPs further towards the mesodermal or neural lineage (Fig. S5F-P), cells displayed dynamics corresponding to those found *ex vivo* (Fig. 3). Thus, NMPs show variable oscillations in Hes1 expression levels that are dependent on Notch signalling.

**Figure 5.**
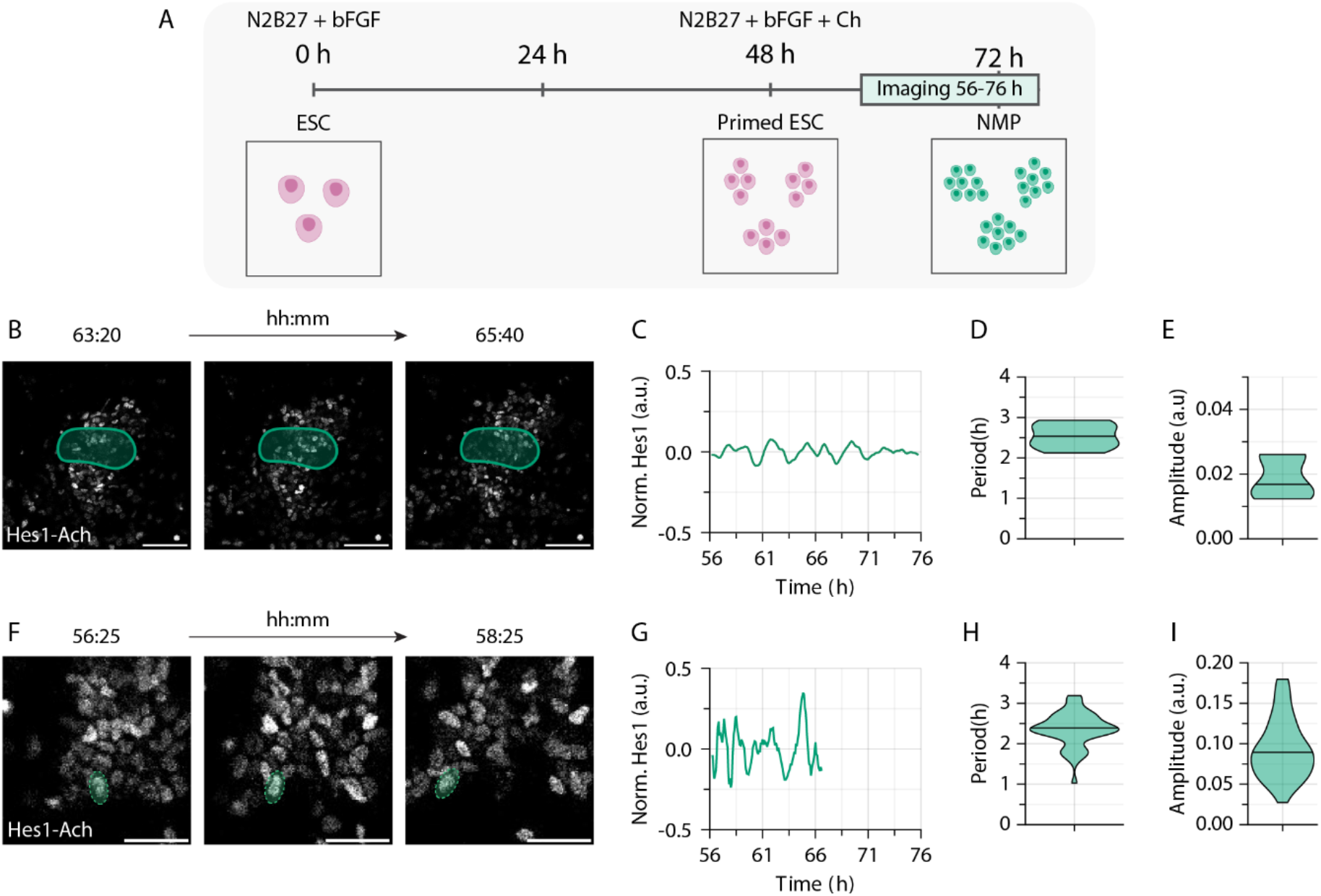
Hes1 shows noisy oscillations in *in vitro* differentiated NMPs. A. Mouse ESCs were differentiated *in vitro* to NMPs. Hes1-Achilles intensity was measured by fluorescence real-time imaging from 56 - 76 h of differentiation. **B-E** Quantification of Hes1-Achilles dynamics in whole NMP colonies (n=8). Representative snapshots are shown in **B**. Timeseries data (mean-normalised) of representative NMP colony is shown in **C**. **D, E** Quantification of the period (**D**) and amplitude (**E**) by wavelet transform. Scale bar 100 µm. **F-I** Quantification of Hes1-Achilles dynamics in single NMP cells (n=46). Single cells were tracked manually to quantify Hes1-Achilles dynamics. Representative snapshots are shown in **F**. Representative timeseries data (mean-normalised) of NMP cells is shown in **G**. **H, I** Quantification of the period (**H**) and amplitude (**I**) by wavelet transform. Scale bar 50 µm. Note that data corresponds to control data in Fig. S5B-E.

### Induced differentiation to neural tube by FGF inhibition results in Hes1 dynamics with increased amplitude in the pre-NT

Previous studies have highlighted the function of Hes dynamics in the anterior NT^6,13,28,29,32,53^. Therefore, we asked how Hes1 dynamics changed upon induced differentiation. As FGF inhibition induces differentiation of pre-NT to NT^25,27,54^ and PSM to somite^55,56^, we inhibited FGF activity in 2D spread-out cultures using SU5402 (Fig. 6). As expected, FGF inhibition led to a rapid regression of the oscillating field in the PSM, which indicated induced differentiation and confirmed effectiveness of the inhibitor (Fig. S6A). In agreement with changes of Hes1 expression levels from ‘posterior’ to ‘anterior’ PSM (Fig. 4F), Hes1-Achilles expression levels at population level were increased in the PSM region with a corresponding increase in amplitude, i.e. the relative amplitude was unchanged, while period remained unchanged (Fig. S6B-D). In contrast, Hes1-Achilles expression levels were slightly decreased in the pre-NT region upon FGF inhibition with the period and relative amplitude being unaffected (Fig. S6E-G).

**Figure 6.**
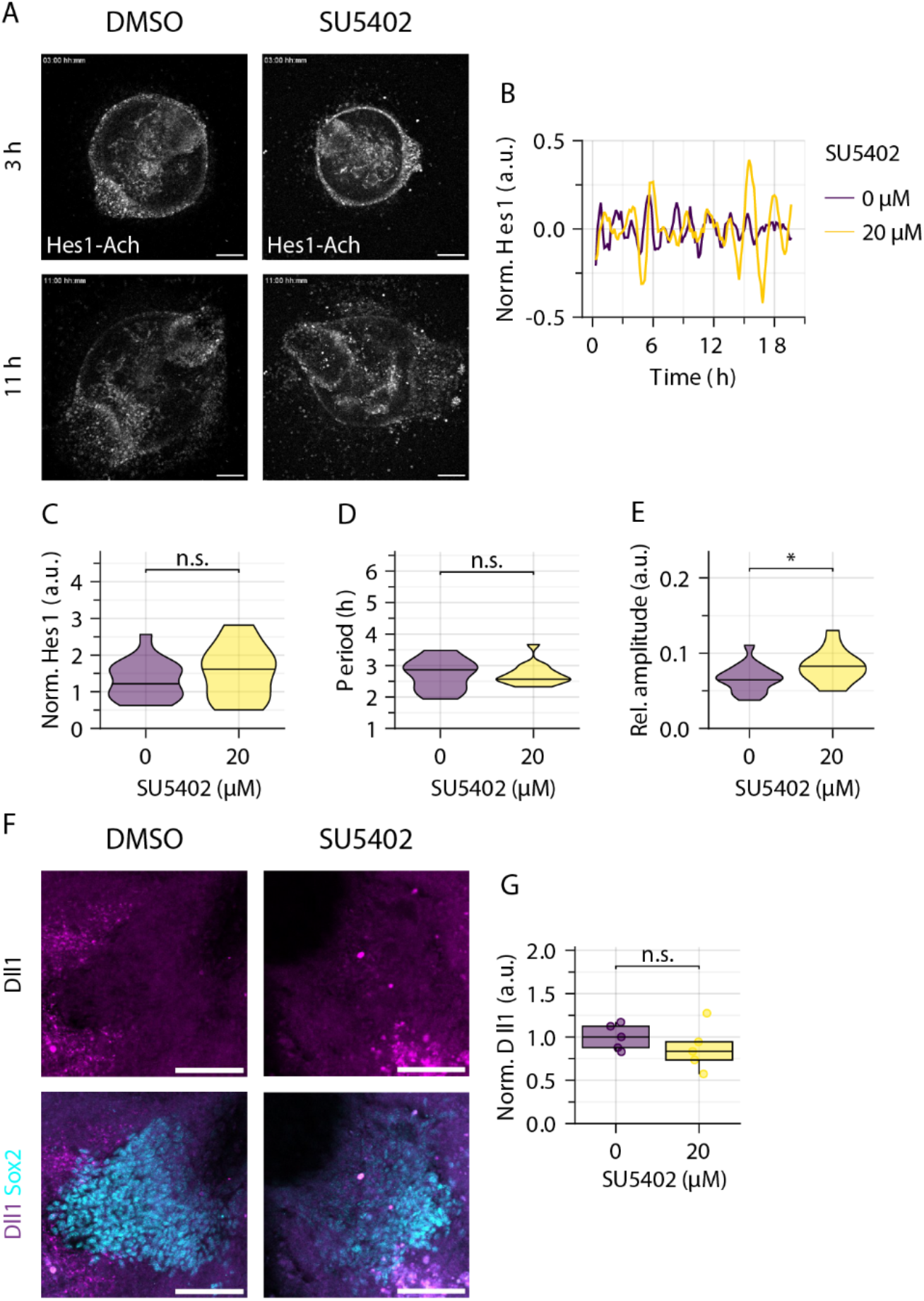
Induced differentiation by FGF inhibition leads to Hes1 dynamics with increased amplitude. Embryonic tail tips were cultured with DMSO control or the FGF receptor inhibitor SU5402. **A** Representative snapshots of the timeseries are shown. Scale bar is 100 µm. **B** Representative timeseries data is shown. **C-E** Quantification of absolute (DMSO-normalised) Hes1-Achilles expression levels (**C**), period (**D**) and relative (mean-normalised) amplitude (**E**) by wavelet transform (0 µM n=15, 20 µM n=15). Further analysis shown in Fig. S6. **F, G** Cultures were fixed after 21 h and immunostaining against Dll1 and Sox2 was performed. Representative images are shown in **F** and the quantification in **G** (0 µM n=5, 20 µM n=5). Dots in boxplots represent individual data points. Scale bar is 50 µm. * is p<0.05, ** is p<0.01, *** is p<0.001, **** is p<0.0001.

To investigate the effect of FGF inhibition on oscillations in single cells, we performed single-cell tracking. In PSM cells, FGF inhibition had a similar effect on Hes1-Achilles dynamics as at population level with increased absolute expression levels and a corresponding increase in amplitude, i.e. the relative amplitude remained unchanged, while the period was maintained (Fig. S6H-J). In contrast, FGF signalling inhibition had a different effect on Hes1-Achilles dynamics in pre-NT cells: absolute expression levels of Hes1-Achilles and the oscillation period remained similar (Fig. 6A-D, S6K, L), whereas the relative amplitude of Hes1-Achilles dynamics increased (Fig. 6E, S6M).

It has been shown recently that expression of the Notch ligand Delta-like 1 (Dll1) begins to oscillate in the NT, adjacent to the last formed somites^11^. To test if differentiation induced by FGF inhibition might have led to changes in Dll1 expression in 2D cultures, we quantified Dll1 by immunostainings. However, we did not detect changes in Dll1 expression levels in the Sox2-positive cell population upon FGF inhibition (Fig. 6F, G, S6N). Thus, this suggests that high FGF signalling levels in pre-NT cells have a dampening effect on Hes1 oscillations and that differentiation leads to an increase in oscillation amplitude, which is not dependent on overall Dll1 protein expression levels.

### Changing Hes1 dynamics by modulating Notch signalling correlates with an increase in cell proliferation

We next addressed if a common mechanism drives Hes1 oscillations in cells of the different tissue types. In general, a delayed negative feedback loop has been suggested to drive such oscillations via repression of Hes genes by Hes proteins themselves^13,24,36,57-60^. In addition to a Hes-driven feedback loop, the Notch ligand Dll1 is dynamically expressed in the PSM^11,61^, which has been proposed to drive a delayed coupling mechanism of segmentation clock oscillations resulting in kinematic waves in the PSM^39,61-63^. In neural tissue, however, Dll1 oscillations were not observed in the pre-NT, but only in the NT adjacent to the last formed somites, where single cells started to oscillate in a salt-and-pepper like pattern^11^.

To clarify the role of Notch signalling in driving Hes1 oscillations in the embryonic tail, we first analysed the expression of Notch signalling components in the different regions of the embryonic tail, especially the pre-NT. We found Notch ligands Dll1, Dll3 and Dll4 and the Notch receptors to be expressed in the PSM, TB and pre-NT by HCR^33^ (Fig. S7A, B). We next performed immunostainings to visualize ligand protein expression levels. In agreement with previous publications^11^, Dll1 expression levels were elevated in the PSM compared to the pre-NT and staining patterns varied between embryonic tails indicating dynamic expression (Fig. 7A and data not shown). However, Dll1 – just like Dll4 – was not only detected in the PSM but also in the pre-NT and TB, albeit at lower expression levels and without indications of dynamics (Fig. 7A). Together with previous findings on Dll1 oscillations in the embryonic tail^11^, this suggests that Dll1 and Dll4 are present at low expression levels in pre-NT cells but not oscillatory, whereas Dll1 expression levels oscillate in PSM cells (Fig. 7B).

**Figure 7.**
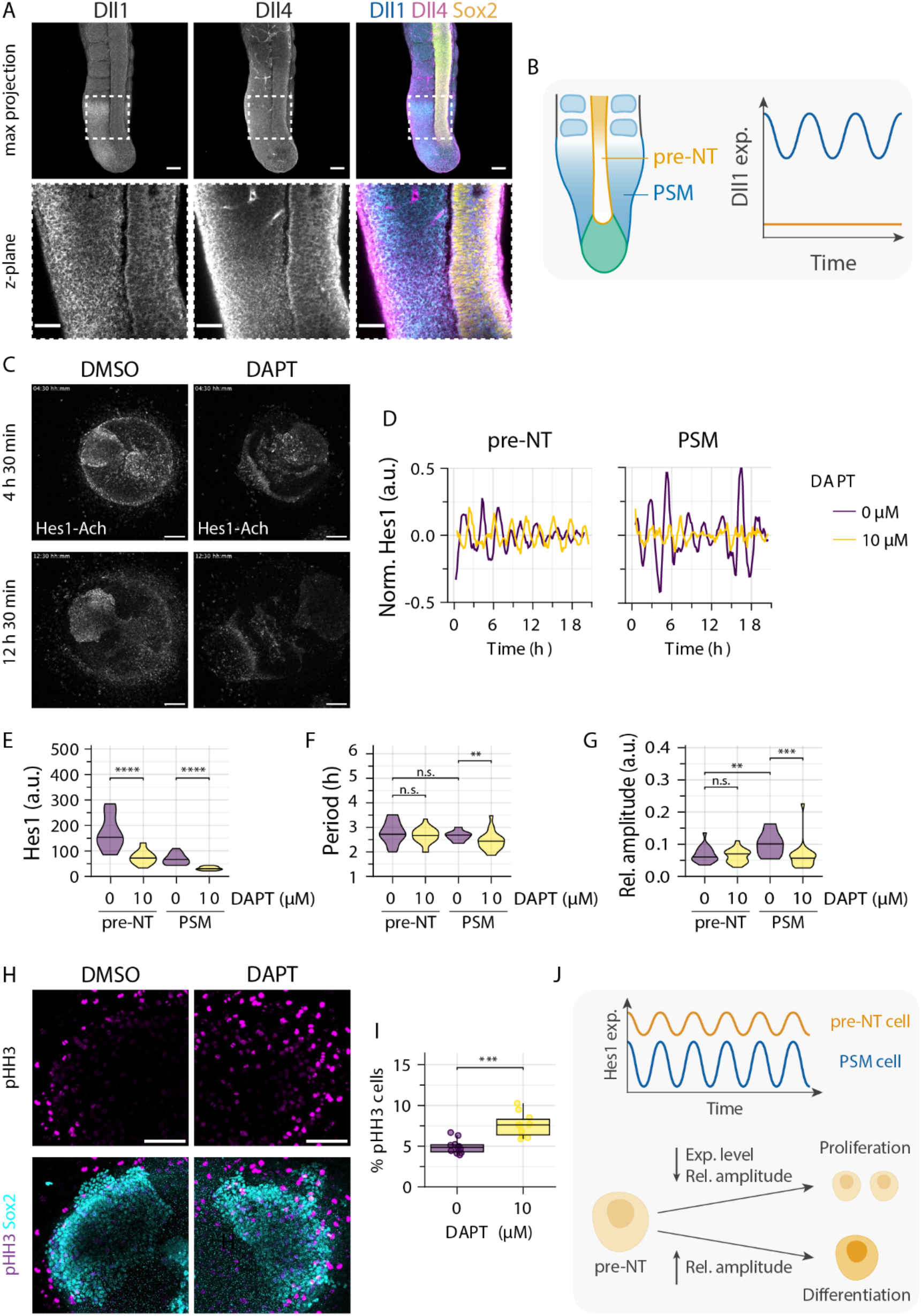
Notch inhibition has a differential effect on Hes1 oscillations in PSM and pre-NT cells. A. Representative image of immunostaining of an E10.5 embryonic tail for Dll1, Dll4 and the neural marker Sox2. Scale bar of top panel is 100 µm and bottom panel is 50 µm. **B** Schematic illustrating that Dll1 oscillates at high levels in the PSM^11^, whereas Dll1 levels are low and non-oscillatory in the pre-NT. **C-I** Embryonic tail tips were cultured with DMSO control or the gamma-secretase inhibitor DAPT. **C** Representative snapshots of the timeseries are shown. Scale bar is 100 µm. **D** Representative timeseries data for pre-NT cells (left panel) and PSM cells (right panel) cells are shown. **E-G** Quantification of absolute Hes1-Achilles expression levels (**E**), period (**F**) and relative (mean-normalised) amplitude (**G**) by wavelet transform. (0 µM n=18, 10 µM n=21). Further quantification shown in Fig. S7. **H, I** Cultures were fixed after 18h and immunostaining against phosphorylated histone H3 (pHH3) and Sox2 was performed. Representative images are shown in **H** and the quantification of pHH3-positive cells in the Sox2-positive region in **I** (0 µM n=11, 10 µM n=10). Dots in boxplots represent individual data points. Scale bar is 50 µm. **J** Schematic summarising lower Hes1-Achilles expression level and higher Hes1-Achilles relative amplitude in PSM cells compared to pre-NT. In addition, decreased Hes1-Achilles expression level and relative amplitude led to proliferation of pre-NT cells, while increased Hes1-Achilles amplitude correlates with differentiation. * is p<0.05, ** is p<0.01, *** is p<0.001, **** is p<0.0001.

To compare the influence of Notch signalling on Hes1 protein dynamics in PSM and pre-NT cells, we inhibited Notch signalling with different doses of DAPT and quantified Hes1-Achilles dynamics within PSM and pre-NT at population level and in single cells (Fig. 7, S7). Analysing dynamics in PSM at population level, we found that Hes1-Achilles expression levels decreased to a similar extent for all DAPT concentrations used (Fig. S7C). The oscillation period was slightly increased at 1 µM DAPT, whereas the relative amplitude was strongly reduced independent of the DAPT concentration used (Fig. S7D, E). This indicates that at population level, Notch inhibition leads to a loss of Hes1-Achilles oscillations in the PSM even at low concentrations, which could be consistent with a loss in synchrony between neighbouring cells^19^. In contrast, in pre-NT (Fig. S7F-H) Notch inhibition led to dose-dependent decrease in Hes1-Achilles expression levels (Fig. S7F) and relative amplitude (Fig. S7H), while no significant change in period was observed (Fig. S7G).

To clarify how Notch inhibition affects oscillations in single cells, we performed single-cell tracking and quantified Hes1-Achilles expression levels in pre-NT in the different conditions. Also at single-cell level, Notch inhibition led to a dose-dependent decrease of Hes1-Achilles expression levels in pre-NT cells (Fig. 7C-E, S7I-K). This correlated with a corresponding decrease in oscillation amplitude with the relative amplitude remaining unchanged (Fig. 7G, S7K). Furthermore, no significant change in period was observed (Fig. 7F, S7J). Conversely, in PSM cells Notch inhibition led to a decrease in both Hes1-Achilles expression levels and the relative amplitude (Fig. 7C-G, S7L-N). Likewise, the Hes1-Achilles oscillation period was decreased with high DAPT concentration (Fig. 7F). Thus, Notch inhibition has differential effects on Hes1 oscillations in the PSM and pre-NT, which implies that the mechanism driving Hes1 dynamics differs between the two cell types.

While the essential function of Notch signalling in somitogenesis has been studied extensively^64-67^, its role in the pre-NT especially with regards to signalling dynamics is less clear, even though it has been implicated in proliferation control^26^. In other regions of the nervous system, Hes1 oscillations also regulate proliferation with high, sustained Hes1 expression levels promoting quiescence^31,32,68^. We therefore addressed how Notch signalling inhibition and a corresponding modulation of Hes1-Achilles dynamics affected pre-NT proliferation. Interestingly, when quantifying the percentage of phospho-histone H3-positive mitotic cells (mitotic index) in the Sox2-positive region, we found that Notch inhibition led to an increase in the mitotic index (Fig 7F, G, S7O). This suggests that lowering Notch signalling and thereby Hes1 expression levels and oscillation amplitude, while leaving the period unchanged, promotes proliferation in the pre-NT.

## Discussion

Here, we have addressed how Hes dynamics change in the context of the developing embryonic tail, in particular focussing on the tissues derived from NMPs. To enable this, we have generated a new and endogenous Hes1 reporter mouse line. It contains the fast-maturing fluorescent protein Achilles, which is placed C-terminally to the Hes1 gene. Importantly, the line is homozygously viable and fertile and allows the investigation of Hes1 dynamics at single-cell resolution in various tissue types (Sonnen, unpublished). Using this new reporter, we compared Hes1 dynamics in the PSM and pre-NT at population and single-cell level.

To make this feasible, we had to optimize the embryonic tail culture protocol to ensure survival of NT cells during *ex vivo* culture. In contrast to the minimal DMEM-F-12 medium, which is optimal for PSM growth and differentiation^69^, we made use of neurobasal medium that is optimized for culture of neural tissues. The latter is not only supplemented with additional nutrients and vitamins, but also contains proteins such as insulin. Which of these components are required for NT and pre-NT survival must be addressed in future studies. While this medium allows NT survival, we do observe morphological defects in somite formation, which are presumably due to the presence of retinoic acid in the medium. Despite these morphological defects, dynamics of the segmentation clock are maintained. This medium therefore allowed us to culture embryonic tails *ex vivo* for the quantification of signalling dynamics in both tissue types in the same culture.

### Differential mechanisms driving Hes1 dynamics in the PSM and pre-NT

In pre-NT cells, Hes1 oscillated at high expression levels with low amplitude, whereas in PSM cells, it oscillated at low expression levels with high amplitude (Fig. 7H). Notch inhibition led to a decrease in Hes1 expression levels and a proportional decrease in the amplitude in pre-NT cells. In contrast, in PSM cells an over-proportional decrease in amplitude was induced, i.e. the relative amplitude decreased, both when analysing Hes dynamics at population and single-cell level. This implies that different mechanisms drive Hes1 dynamics in the two tissue types. While the effect of Notch inhibition on Hes1 dynamics in pre-NT cells is consistent with a delayed negative feedback loop downstream of constant Notch signalling^57,58^, oscillations in PSM cells appear to be further amplified in a Notch-dependent manner.

In fact, the Notch ligand Dll1 has been shown to oscillate in various cell types including the PSM, which results in periodic activation of proneural gene expression^11,70,71^. Previously, Dll1 oscillations were not detected in the pre-NT, but only in the NT adjacent to the somites^11^. We have shown here that even in the pre-NT both Dll1 and Dll4 are present at low expression levels. Interestingly, in a study using cultured cancer cell lines it was shown that Dll1 expression induces dynamic expression, while Dll4 leads to more sustained expression of downstream target genes^72^. How the presence of each ligand and the dynamics of Dll1 exactly influence Hes1 dynamics in PSM and pre-NT cells has to be addressed in future studies.

In the PSM, oscillatory Dll1 expression has been suggested to lead to kinematic waves travelling through the PSM^11,63,73^. Loss of Notch-dependant intercellular coupling could therefore explain the loss of population-wide dynamics we observed already at low Notch inhibitor concentrations. However, the oscillation amplitude was also reduced in single-cell oscillations upon Notch inhibition arguing against a sole effect on synchronicity between oscillators. The fact that high Notch inhibitor levels also led to a decrease in the relative Hes1 oscillation amplitude in the pre-NT at population level suggests that Notch signalling might induce some form of synchronization between pre-NT cells, as suggested recently for Hes5 in the NT^74^. While our mosaic labelling approach for single-cell tracking allows us to track cells and their offspring, it does not allow the quantification of Hes1 dynamics in all neighbours of a given cell. Therefore, the intercellular synchronization between single-cell oscillators has to be addressed in future investigations.

### Implications for the role of Hes1 dynamics in the preneural tube

Hes1 dynamics have been found to regulate proliferation and differentiation in the anterior developing NT^13,68,75^. In undifferentiated cells, Hes1 oscillates in an alternating fashion with proneural genes, while Hes1 is switched off in differentiating cells. Moreover, high absolute expression levels regulate the switch between quiescence and proliferation in adult stem cells with high expression levels maintaining a quiescent state^31,32,68^. It is therefore interesting to note that we find Hes1 expression levels highest in the pre-NT and expression decreases towards the NT. When inducing a decrease in Hes1 expression levels and oscillation amplitude by inhibiting Notch signalling, the mitotic index was indeed increased. Similar to morphogen gradients in the PSM, FGF and Wnt signalling are high in the pre-NT. High FGF signalling prevents differentiation of the pre-NT to NT^25,27,54-56^. By quantifying Hes1 dynamics in pre-NT cells upon inhibition of FGF signalling, we found that the relative amplitude of Hes1 dynamics was increased (Fig. 7H). This indicates that FGF has a dampening effect on Hes1 dynamics in the pre-NT and that reduced FGF signalling results in Hes1 dynamics with higher relative amplitude. Whether FGF signalling directly affects Hes1 dynamics or functions indirectly via induction of differentiation has to be disentangled in the future.

Together, this supports a hypothesis in which cells in the pre-NT are maintained in an undifferentiated, anticipating phase until they become part of the NT, where cells differentiate further in conjunction with the neighbouring somites.

### Information transmission by noisy Hes1 oscillations

We have quantified Hes1 oscillations in single PSM, NMP and pre-NT cells. In both NMP and pre-NT cells, Hes1 oscillations are noisy at high expression level with low amplitude. In contrast, in the PSM single-cell oscillations are less noisy at low expression level with higher amplitude (Fig. 7H). In addition to these single-cell oscillations, it is well known that oscillations between neighbouring PSM cells are coupled leading to travelling waves of signalling along the tissue^41,47,76^. These coupled, highly synchronized oscillations ensure that information for proper segmentation can be transmitted from posterior to anterior in every segmentation cycle.

Even though the pre-NT lies directly adjacent to this oscillating field, the tissues are spatially separated by epithelialization and extracellular material. In the TB and pre-NT, we detect different expression patterns of several signalling components and downstream targets of the Wnt and Notch signalling pathway compared to the PSM (this study and MAMEP database). In correlation with this, Hes1 dynamics in pre-NT cells and NMPs are not only noisy but also less synchronized between neighbouring cells. The absence of synchronized oscillations or travelling waves implies that information in the pre-NT is not transmitted along the anterior-posterior axis via these dynamics. Conversely, Hes1 presumably has a more local function in regulating cellular behaviour on a single-cell basis, as has been suggested in other tissues^70,71^.

Notably, once the pre-NT has differentiated to the NT, it has been shown that signalling between somite and NT does regulate further patterning and segmentation of the developing NT in conjunction with the differentiating somite^77^. This ensures that spinal cord, nerves and vertebrae form one unit. Thus, proper patterning of the embryo’s spinal axis would be dependent on proper somitogenesis. Based on this, somitogenesis would therefore have to be tightly regulated and coupled to axial elongation, which necessitates highly accurate information transmission. In contrast, differentiation and segmentation of the NT would be downstream of somite formation, which suggests that information could be kept local with the sole aim to allow proliferation and prevent faulty or premature differentiation of single cells in the pre-NT.

In summary, Notch-dependent Hes1 dynamics in single cells of the developing embryonic tail differ in expression levels, amplitude and noise, which has implications for information transmission in the PSM and NT. The detailed molecular mechanism of how Hes dynamics are generated and what information is stored in these dynamics in different cell types during embryonic development are important questions for future studies.

## Acknowledgements

We thank all members of the Sonnen, Bothma and Garaycoechea labs for feedback and discussions. We are thankful to Juan Garaycoechea and Jacques Bothma for feedback on the manuscript. We thank Jan-Daniël de Leede for help with data analysis, Valerie Wilson, Mina Gouti and Jana Wolf for useful discussions and Andreas Sonnen for help with Alphafold. We thank Jeroen Korving, Stieneke van den Brink, Joris Geigenmüller, Tjerk Swinkels and Mariska Dijkers for technical help. This work was supported by the Hubrecht imaging and animal facilities. The H2B-mCherry was kindly provided by the Riken Center for Biosystems Dynamics Research. The T-Cre/ERT2, RosaCreERT and PGK-Cre lines were kindly provided by Jackson Laboratories. The LuVeLu reporter line was kindly provided by Olivier Pourquié, the Achilles-Hes7 by Ryoichiro Kageyama and the Achilles construct by Atsushi Miyawaki (Riken Center for Brain Science). P.S. was supported by KWF grant 13661 from the Dutch Cancer Foundation. This work was supported by the Hubrecht Institute and received funding from the European Research Council under an ERC starting grant agreement no. 850554 to K.F.S. In addition, this research was supported in part by grant NSF PHY-1748958 to the Kavli Institute for Theoretical Physics (KITP).

## Author contributions

Conceptualization: Y.A., P.S. and K.F.S., Methodology: Y.A., Software: P.S., Formal Analysis: P.S., Y.A. and K.F.S., Investigation: Y.A., M.J.O, W.T. and K.F.S., Resources: W.T. and W.H.M.M., Writing – Original draft: Y.A. and K.F.S., Writing – Review and Editing: Y.A., K.F.S., M.B., Supervision, Project Administration and Funding acquisition: K.F.S.

## Star Methods

### Contact for reagent and resource sharing

Further information and requests for resources, data or code should be addressed to the Lead Contact, Katharina Sonnen.

### Experimental model and subject details

### Mouse lines

All animals were housed and bred according to institutional guidelines, and procedures were performed in compliance with Standards for Care and Use of Laboratory Animals with approval from the Hubrecht Institute ethical review board. All animal experiments were approved by the Animal Experimentation Committee (DEC) of the Royal Netherlands Academy of Arts and Sciences.

The Hes1-Achilles knock-in reporter line was generated employing standard gene targeting techniques using IB10 ESCs. To generate Hes1-Achilles alleles, we targeted the stop codon of the endogenous Hes1 locus with a reporter cassette coding for a linked fluorophore (Hes1-linker-Achilles). The reporter cassette is linked to the Hes1 gene via a flexible GSAGS sequence and includes a selection cassette. After generation of the knock-in reporter line, the selection cassette was removed by Cre-mediated excision to yield the final Hes1-Achilles allele, using the PGK-Cre mouse line^78^.

The LuVeLu^47^ and Achilles-Hes7^39^ line were published previously. The R26R-H2B-mCherry^79^ (“H2B-mCherry”) line (http://www.clst.riken.jp/arg/reporter_mice.html) was obtained from the Riken Center for Biosystems Dynamics Research (accession number CDB0204K). The Tg(T-cre/ERT2)1Lwd/J^80^ (“T-Cre/ERT2”) line was obtained from the Jackson Laboratory (accession number #025520). The Tg(Pgk1-cre)1Lni/CrsJ^78^ (“PGK-Cre”) line was obtained from the Jackson Laboratory (accession number #020811). The R26-CreERT2^81^ (“RosaCreERT”) line was obtained from the Jackson Laboratory (accession number #00846).

### Cell lines

For the generation of Hes1-Achilles_RosaCreERT ESCs, E3.5 blastocysts were isolated from pregnant Hes1-Achilles_RosaCreERT females. Each blastocyst was transferred to a single well of a 24-well plate seeded with feeder MEFs. After 6 days, cells were trypsinized and expanded into a 12-well plate with feeders. Four days later, cells were expanded into 6-well plates, and simultaneously used for extraction of genomic DNA followed by polymerase chain reaction (PCR) to determine the genotype. After confirmation of genotype, cells were further expanded for one more passage prior to freezing down. ESCs were cultured in N2B27-2i medium as previously described^82^.

The Sox2-H2B-iRFP (“Sox2-iRFP”) knock-in reporter line was generated employing CRISPaint^51^ gene targeting techniques using Lipofectamine 3000 (ThermoFisher, L3000001) and Hes1-Achilles_RosaCreERT ESCs. To generate Sox2-iRFP alleles, we targeted a PAM site before the stop codon of the endogenous Sox2 locus with a reporter cassette coding for a separated H2B gene linked to a fluorophore (Sox2-T2A-H2B-linker-iRFP). The reporter cassette is separated from the Sox2 gene via a small 2A sequence. Furthermore, the fluorophore is linked to the H2B gene via a flexible GSAGS sequence. The selection cassette is also separated from the reporter cassette via a small 2A sequence. Selection occurred two days after transfection and lasted three days. Colonies were picked and transferred to a single well of a 96-well plate seeded with irradiated feeder MEFs. Colonies were then expanded, genotyped and frozen as explained above.

## Methods details

### Transgenic, mouse strains and animal work

For all experiments (unless stated otherwise) female mice were sacrificed on 10.5 dpc and embryos dissected. Dissections were performed in PBS supplemented with 1 % Bovine Serum Albumin (Biowest, P6154), 200 μg Penicillin-Streptomycin (Gibco, 15140122), 1x GlutaMAX (Gibco, 35050038) and 10 % Glucose Solution 45 (Sigma-Aldrich, G8769).

*Ex vivo* culture was performed as described previously^46,47,69^. The neurobasal medium used to culture *ex vivo* samples consists Special Advanced DMEM/F-12 (Cell Culture Technologies) and Neurobasal medium (Gibco, 12348017) (1:1), 1x N2 (Gibco, 17502048), 1x B27 (Gibco, 12587010), 1x GlutaMAX (Gibco, 35050038), 0.004 % Bovine Serum Albumin (Biowest, P6154), 0.1 mM 2-Mercaptoethanol and 200 μg Penicillin-Streptomycin (Gibco, 15140122) The following small molecule inhibitors dissolved in DMSO were used at the concentration indicated in the text and Figure legends: FGF-basic (“bFGF”, Peprotech, 450-33), CHIR99021 (“Chiron”, Sigma-Aldrich, SML1046), DAPT (Sigma-Aldrich, D5942), IWP-2 (Sigma-Aldrich, I0536), Retinoic acid (“RA”, Sigma-Aldrich, R2625-50MG), and SU5402 (Sigma-Aldrich, SML0443). Samples were imaged in 8-well Chambered Coverglass w/ non-removable wells (ThermoFisher, 155411PK)

For experiments including Hes1-Achilles mice, individual mice were genotyped by PCR (Fig. S2) using primers Hes1-C-forward (GACCTCGGTGGGTCCTAACGC), linker-Achilles-reverse (CTTGCCGGTGGTGCAGATCAG) and Hes1-N-reverse (GAGGTGGGCTAGGGACTTTACGG).

### *In vitro* differentiation

ESCs were differentiated to NMP, mesodermal or neural fate as described previously^50,52^. To minimise fluorescent background, Special Advanced DMEM/F-12 (Cell Culture Technologies), without phenol red was used. Every time, cells were differentiated and imaged in µ-Dish 35 mm Quad (Ibidi, 80416).

### Immunostaining

For immunostaining, samples were fixed in 4 % formaldehyde for 30 minutes and kept in PBS at 4°C. Commercial antibodies were used including anti-Dll1 rat (MABN2284, Sigma-Aldrich), anti-Dll4 goat (AF1389-SP, R&D systems), anti-Cleaved Caspase-3 rabbit (9661S, CST), anti-phospho Histone H3 mouse (05-806, Sigma-Aldrich), Alexa Fluor 555 Phalloidin (A34055, ThermoFisher), anti-Tbx6 rabbit (ab38883, abcam), anti-Sox2 mouse (ab79351, abcam), anti-Sox2 rabbit (ab92494, abcam), anti-T human/mouse (AF2085-SP, R&D systems), anti-Pax6 recombinant (ab195045, abcam), Goat anti-Rat Alexa Fluor 555 (A-21434, ThermoFisher), Donkey anti-Rabbit Alexa Fluor 488 (A-21206, ThermoFisher), Donkey anti-Rabbit Alexa Fluor 568 (A10042 ThermoFisher), Donkey anti-rabbit Alexa Fluor 405 (A48258, ThermoFisher), Donkey anti-mouse Alexa Fluor 647 (A-31571, ThermoFisher), Donkey anti-rat Alexa Fluor 555 (A78945, ThermoFisher), Donkey anti-goat Alexa Fluor 647 (A32849, ThermoFisher), Donkey anti-mouse Alexa Fluor 568 (A10037, ThermoFisher) and Donkey anti-mouse Alexa Fluor 647(A-31571, ThermoFisher). We stained samples with primary and secondary antibodies overnight at a 1:1000 dilution in PBST (0.1 % Tween™ 20) containing 1 % Bovine Serum Albumin (Biowest, P6154).

### *In situ* hybridization and hybridization chain reaction

Probe generation and *in situ* hybridization were described previously^18^. Probes against Axin2, Dusp4, Hes7 and Lfng were used as described in the literature^18,83,84^. Probes against Achilles and Hes1 were generated using the full-length cDNA^85^. Subsequently samples were dehydrated, paraffin embedded, and sectioned. Standard H&E staining was performed.

Hybridization chain reaction was performed as described previously^33^. Probe sets for the following gene targets: Dll1, Dll3, Dll4, Hes5, Hes7, Meox1, Notch1, Notch2, Notch3, Notch4 and Pax6 were generated^86^. All animals for HCR were stained once using the same hybridization, washing and amplification time for all gene targets.

Images of ISHs were taken with a Leica M165 FC stereomicroscope using 8:1 zoom. Images of HCR were taken with a Leica TCS SP8 MP confocal microscope using a 10x and 20x objective (details below). Brightness and contrast were adjusted uniformly to the entire image.

### Confocal microscopy

Imaging was performed using a Leica TCS SP8 MP confocal microscope featuring an incubator and gas mixer for CO_2_, O_2_ and temperature control. Samples were excited with a OPSL Laser at a wavelength of 514 nm (Hes1-Achilles, Achilles-Hes7 and LuVeLu) or 568 nm (H2B-mCherry) or 638 nm (Sox2-iRFP) through a HC PL APO CS2 20x/0.75 DRY objective. For *ex vivo* culture experiments, a z-stack of 10-12 planes at 10 µm distance was scanned every 10 min. For *in vitro* differentiation experiments, a z-stack of 3-4 planes at 5 µm distance was scanned every 5 min. Multiple samples were recorded using a motorized stage during each experiment. Movies were recorded in 512 x 512 pixels (pixel size 1,517 µm). For imaging of immunostaining and HCR samples, a z-stack of 10-12 planes at 10 µm distance was scanned in 1024 x 1024 pixels (pixel size 0,758 µm).

## QUANTIFICATION AND STATISTICAL ANALYSIS

### Image and data processing

For quantification of mean intensity or number of positive cells in fixed sample imaging, Fiji^87^ was used. Kymographs were generated as described previously^69^. For quantification of mean intensity over time, ROImanager, Mastodon or manual tracking were used. Quantification of oscillation dynamics were performed using pyBOAT^49^ and Julia. For determination of main period and amplitude, sample tracks were averaged and normalised (either by mean or DMSO, as stated in the legends) using Julia. Furthermore, all graphs and heatmaps are plotted using Julia.

### Statistical analysis

For all experiments at least three independent experiments were performed. To visualize data, Tukey style boxplots were used (bar=median, box=25th and 75th percentile, whiskers=1.5*IQR). Data displayed as violin plots present data from max to min with bar=median. For statistical comparisons between groups, Mann-Whitney U tests were computed in Julia.

## DATA AND SOFTWARE AVAILABILITY

Julia scripts are available upon request.

## KEY RESOURCES TABLE

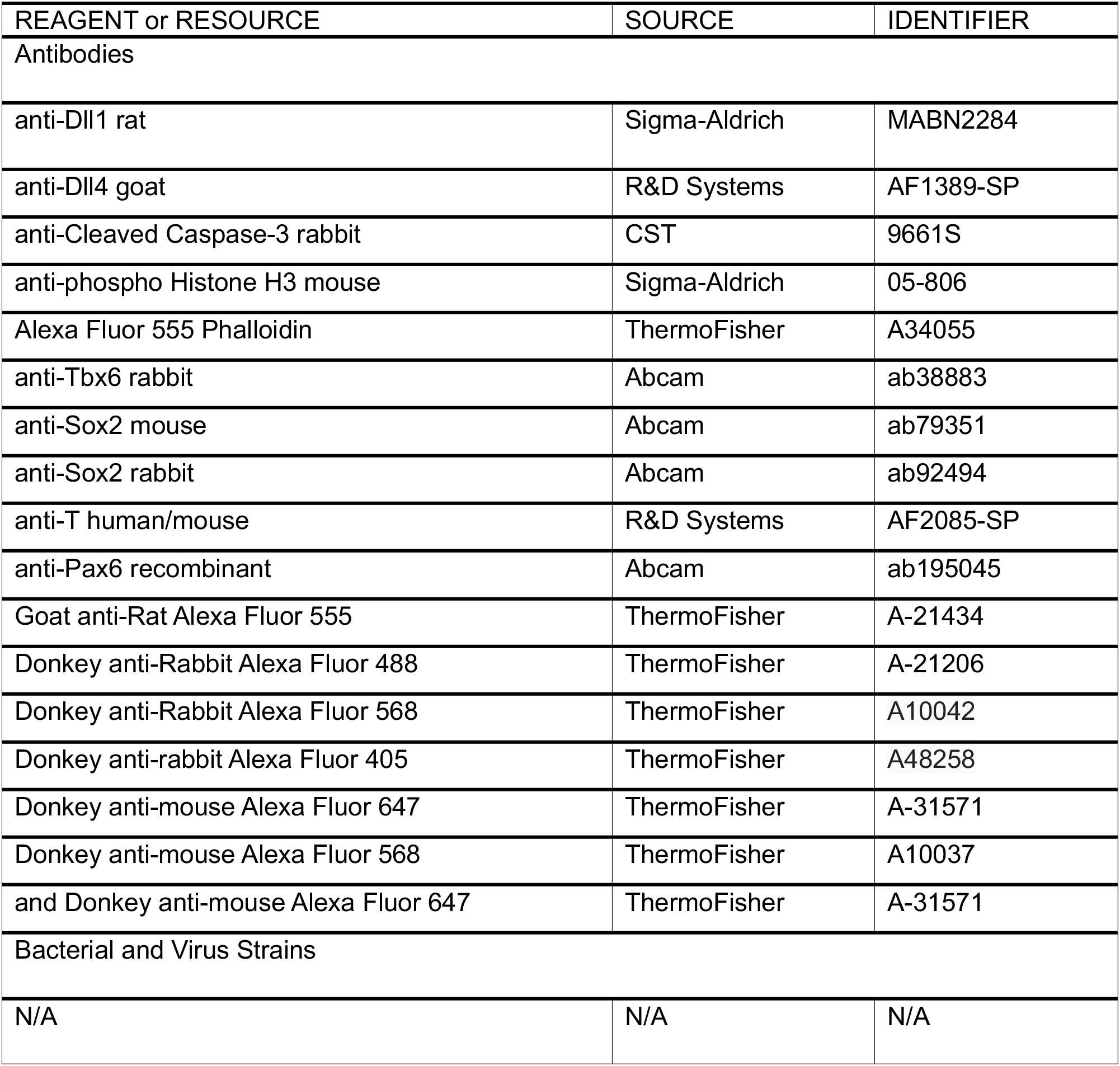
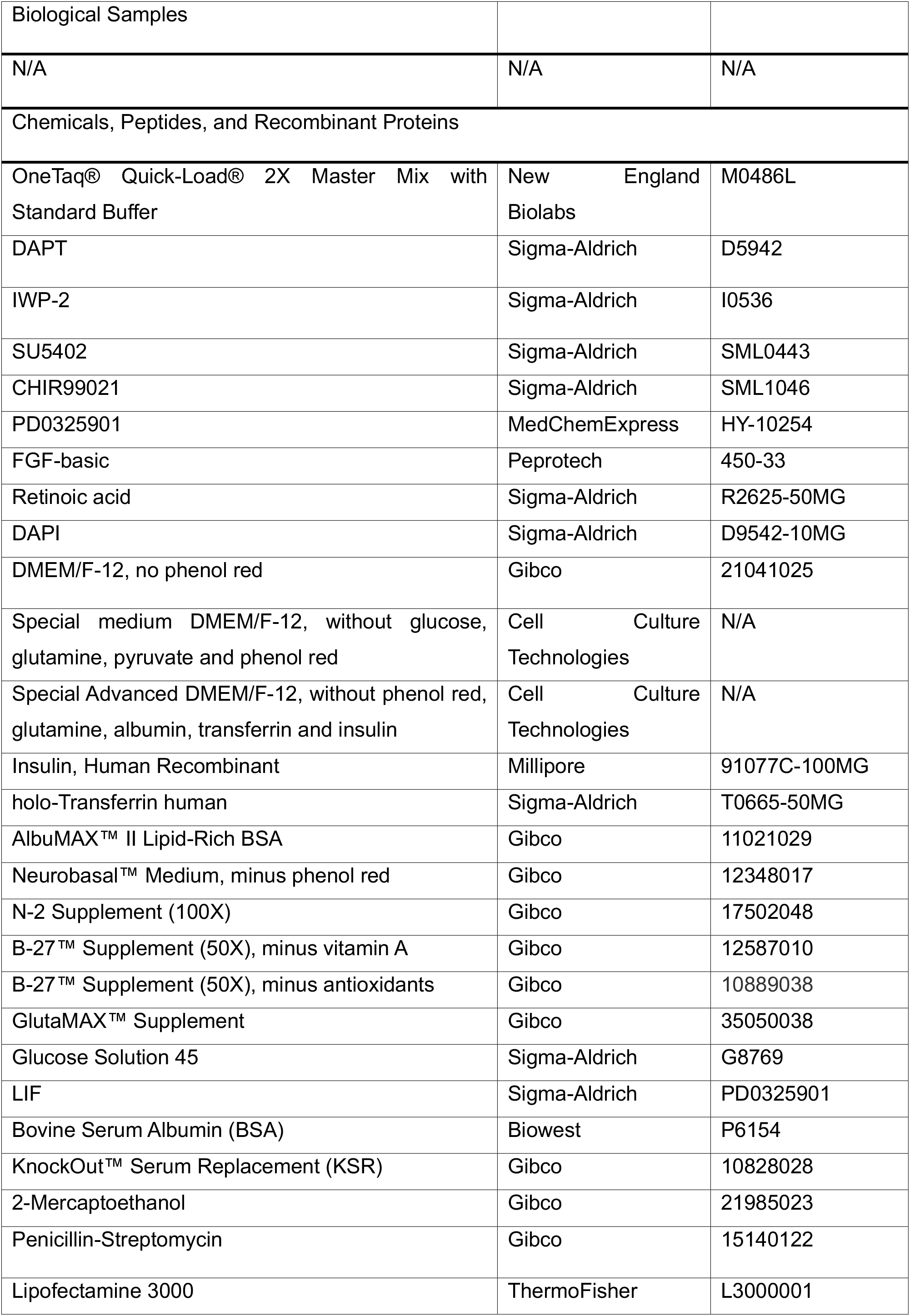
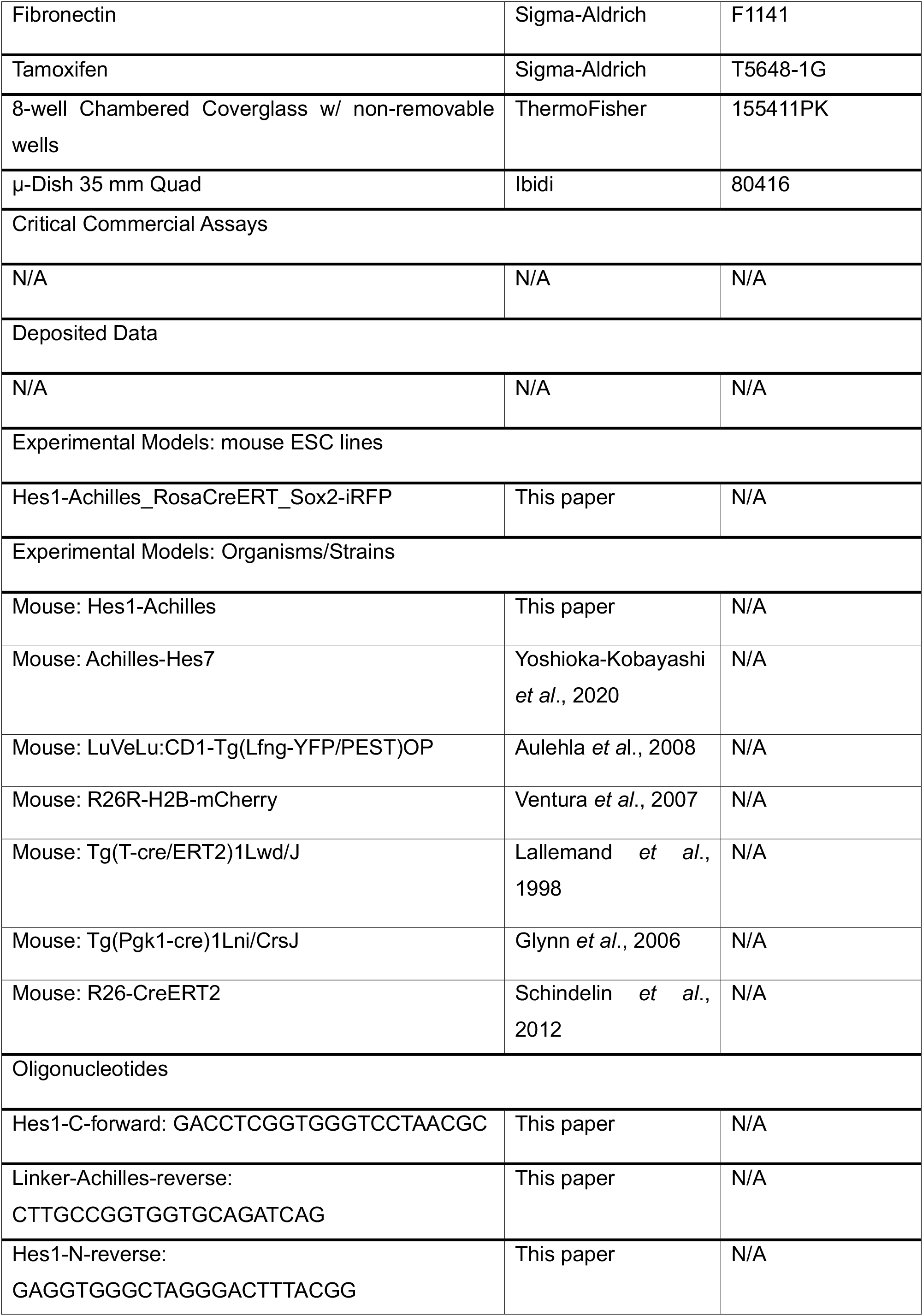
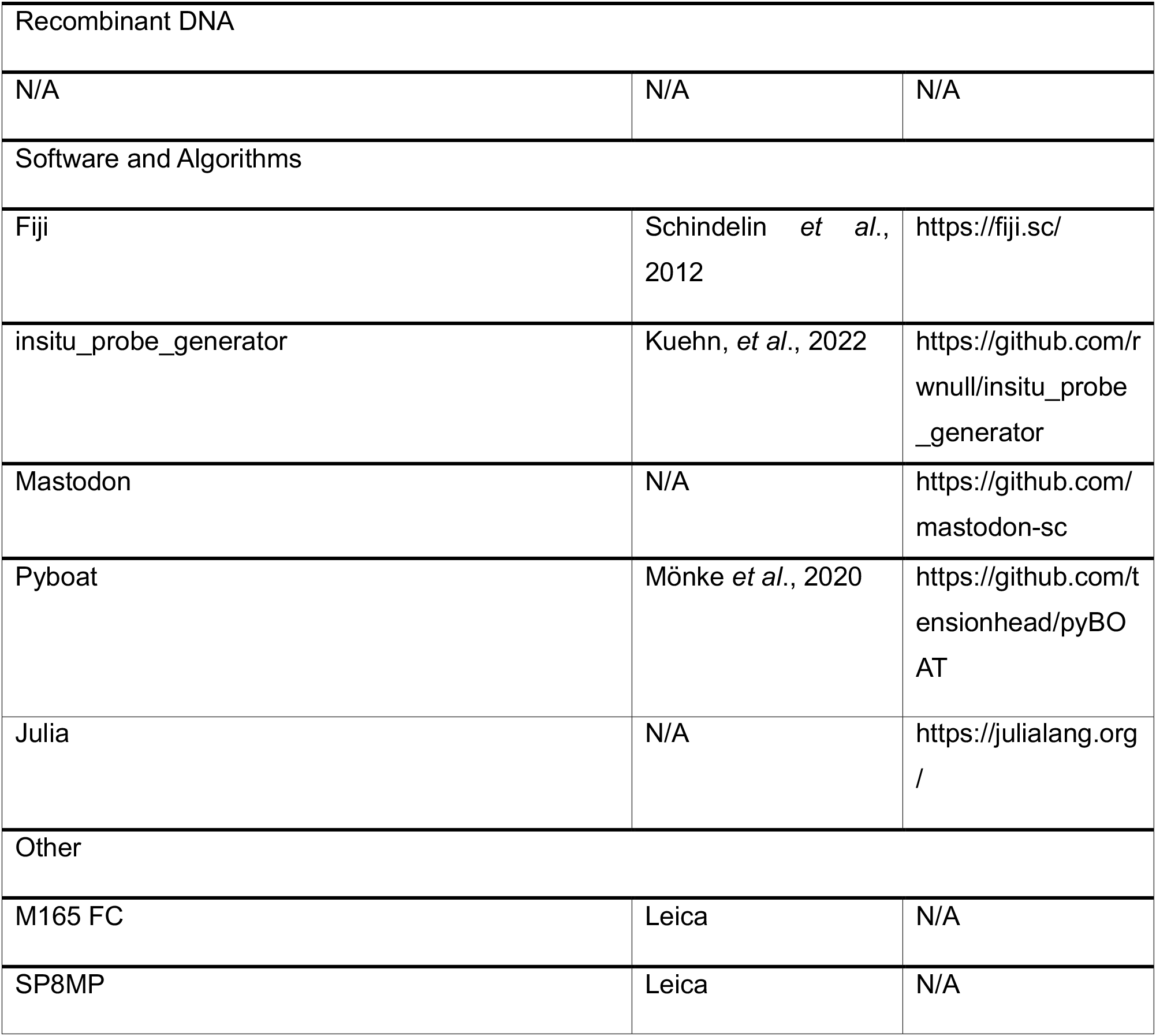

## Supplemental Information

The supplemental material contains 7 supplemental figures, 2 movies and 1 file containing timeseries data.

---

**Figure S1.**
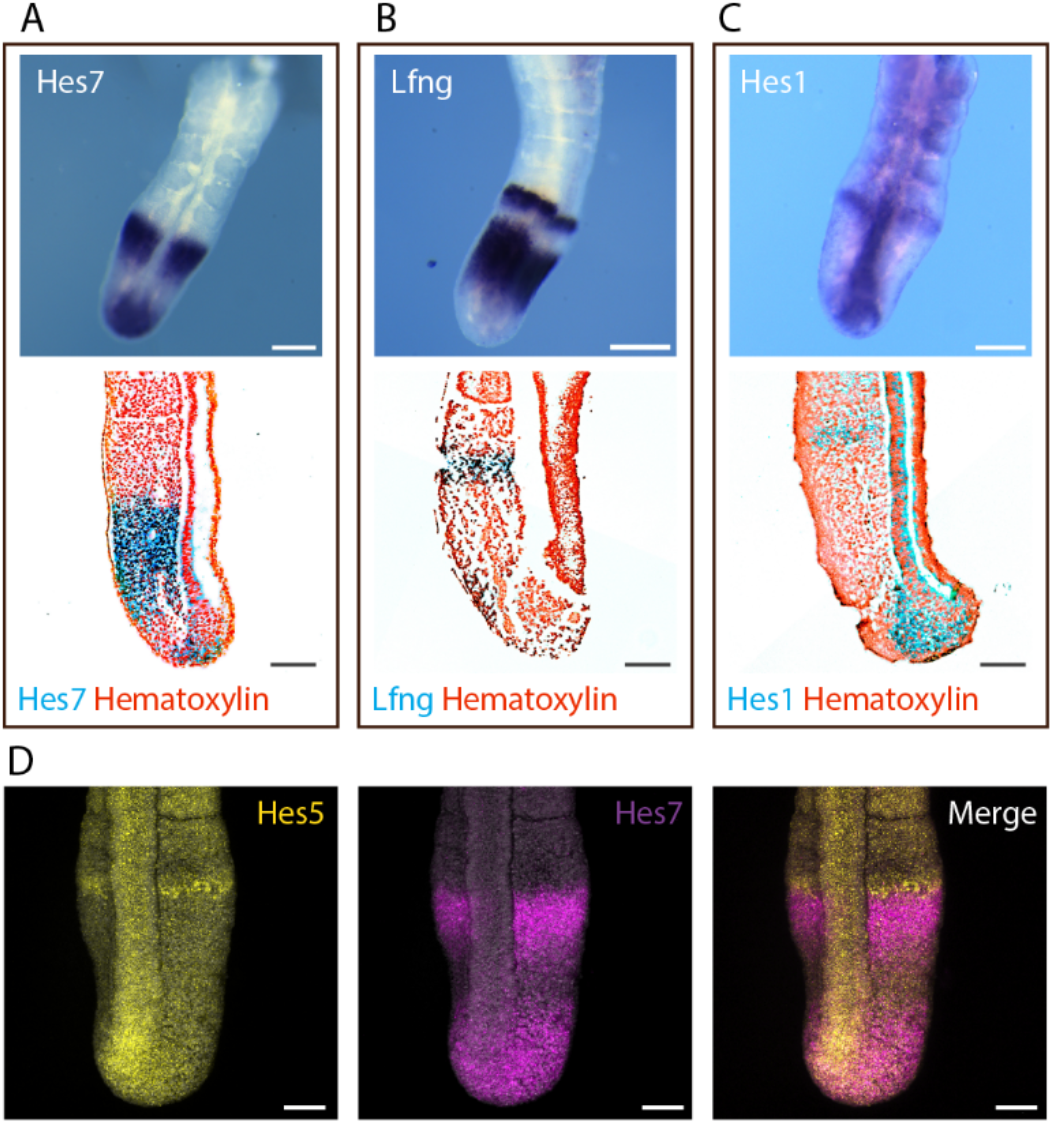
Hes1 and Hes5 are expressed in the pre-NT and TB region of the E10.5 embryo. Visualization of mRNA levels in E10.5 embryonic tails by ISH (**A-C**) and HCR (**D**). **A-C** ISH of Hes7 (**A**), Lfng (**B**) and Hes1 (**C**) (**C** as shown in Fig. 1C, F). Upper panel shows representative image of whole embryonic tail, lower panel 5 µm section counterstained with Hematoxylin. Note the high expression levels of Hes1 in the pre-NT. **D** Expression of target genes Hes5 and Hes7 in embryonic tails visualized by HCR. Scale bar is 100 µm.

**Figure S2.**
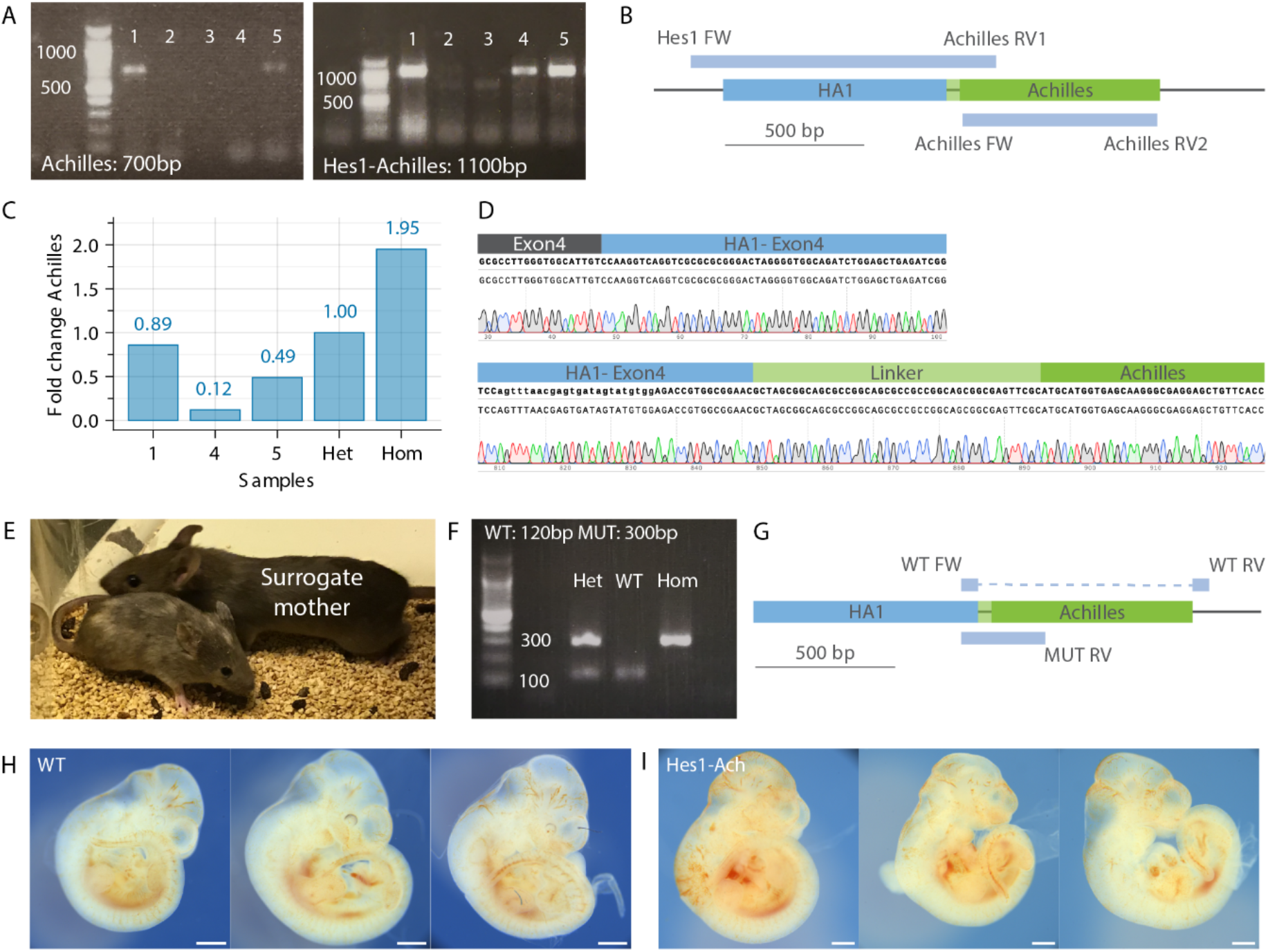
Characterization of Hes1-Achilles reporter mouse line. **A, B** Genotyping for positive clones using primers (forward: FW; reverse: RV) against Achilles and against a product that crosses the homology arm (HA). In **A** a representative gel and in **B** a scheme of the expected gene locus are shown. **C** Integration of exactly one copy of Achilles was confirmed by qPCR (heterozygous: Het; homozygous: Hom). **D** Example of sequencing result of a correctly integrated clone. **E** Chimeric mice were generated by blastocyst injection. **F, G** Genotyping of positive mice is performed using primers to detect integration wildtype (WT) and mutated (MUT) alleles. In **F** representative gels and in **G** a scheme of the expected gene locus are shown. **H, I** Representative images of E10.5 wildtype (WT) (**H**) and Hes1-Achilles homozygous embryos (**I**) are represented. Scale bar is 500 µm.

**Figure S3.**
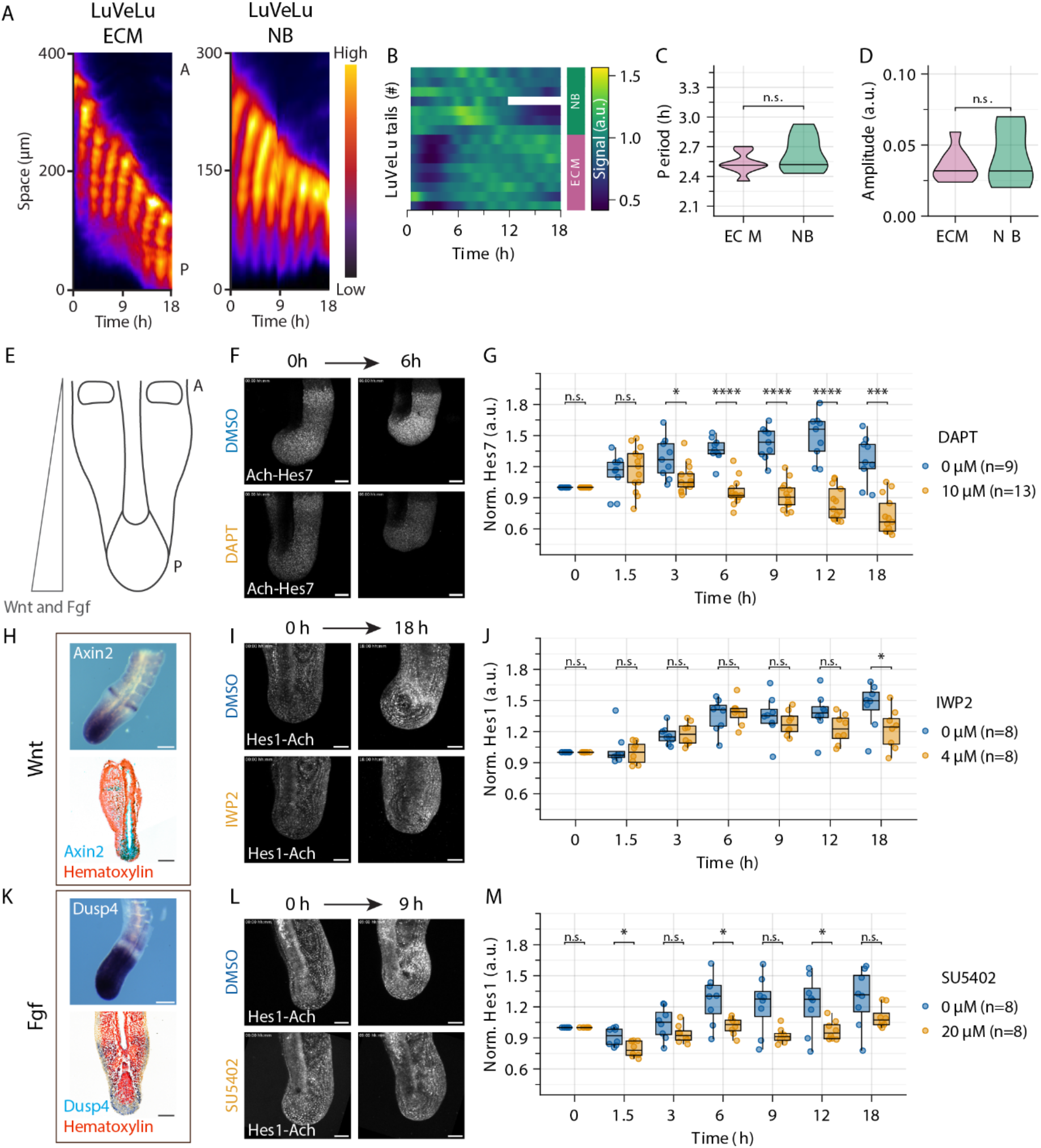
Differential effect of signalling perturbation on *ex vivo* cultured E10.5 embryonic tails. **A-D** Embryonic tails were cultured *ex vivo* in either ECM or NB medium. Intensity of the dynamic signalling reporter LuVeLu was measured. **A** Kymographs generated along the PSM from the posterior (P) to the anterior (A) of a growing embryonic tail cultured in embryo culture medium (ECM, left panel) or neurobasal (NB) medium (right panel) were generated. Note the wave dynamics visible in both. **B** Heatmap of samples illustrating signalling dynamics in the different culture conditions using LuVeLu as reporter line. **C, D** Quantification of period (**C**) and amplitude (**D**) of the signalling dynamics shown in **B** (ECM: n=8 tails; NB: n=7 tails) **E-M** Effect of signalling inhibition on *ex vivo* cultured embryonic tails. **E** Schematic representation of embryonic tail highlighting the gradients of Wnt and FGF along the tissue. **F, G** E10.5 embryonic tails were cultured *ex vivo* and incubated with DMSO control or the gamma-secretase inhibitor DAPT. Achilles-Hes7 intensity was measured over time. In **F** representative images and in **G** the quantification of Hes1-Achilles intensity (normalised to the first timepoint in the embryonic tail are shown. **H** ISH of E10.5 embryonic tail for Wnt target gene Axin2. Upper panel shows representative image of whole embryonic tail, lower panel a 5 µm section counterstained with Hematoxylin. **I, J** E10.5 embryonic tails were cultured *ex vivo* and incubated with DMSO control or the porcupine inhibitor IWP2. Hes1-Achilles intensity was measured over time. In **I** representative images and in **J** the quantification of Hes1-Achilles intensity (normalised to the first timepoint) in the embryonic tail are shown. **K** ISH of E10.5 embryonic tail for FGF target gene Dusp4. Upper panel shows representative image of whole embryonic tail, lower panel a 5 µm section counterstained with Hematoxylin. **L, M** E10.5 embryonic tails were cultured *ex vivo* and incubated with DMSO control or the FGF receptor inhibitor SU5402. Hes1-Achilles intensity was measured over time. In **L** representative images and in **M** the quantification of Hes1-Achilles intensity (normalised to the first timepoint) in the embryonic tail are shown. Scale bar is 100 µm. Dots in boxplots represent individual data points. * is p<0.05, ** is p<0.01, *** is p<0.001, **** is p<0.0001.

**Figure S4.**
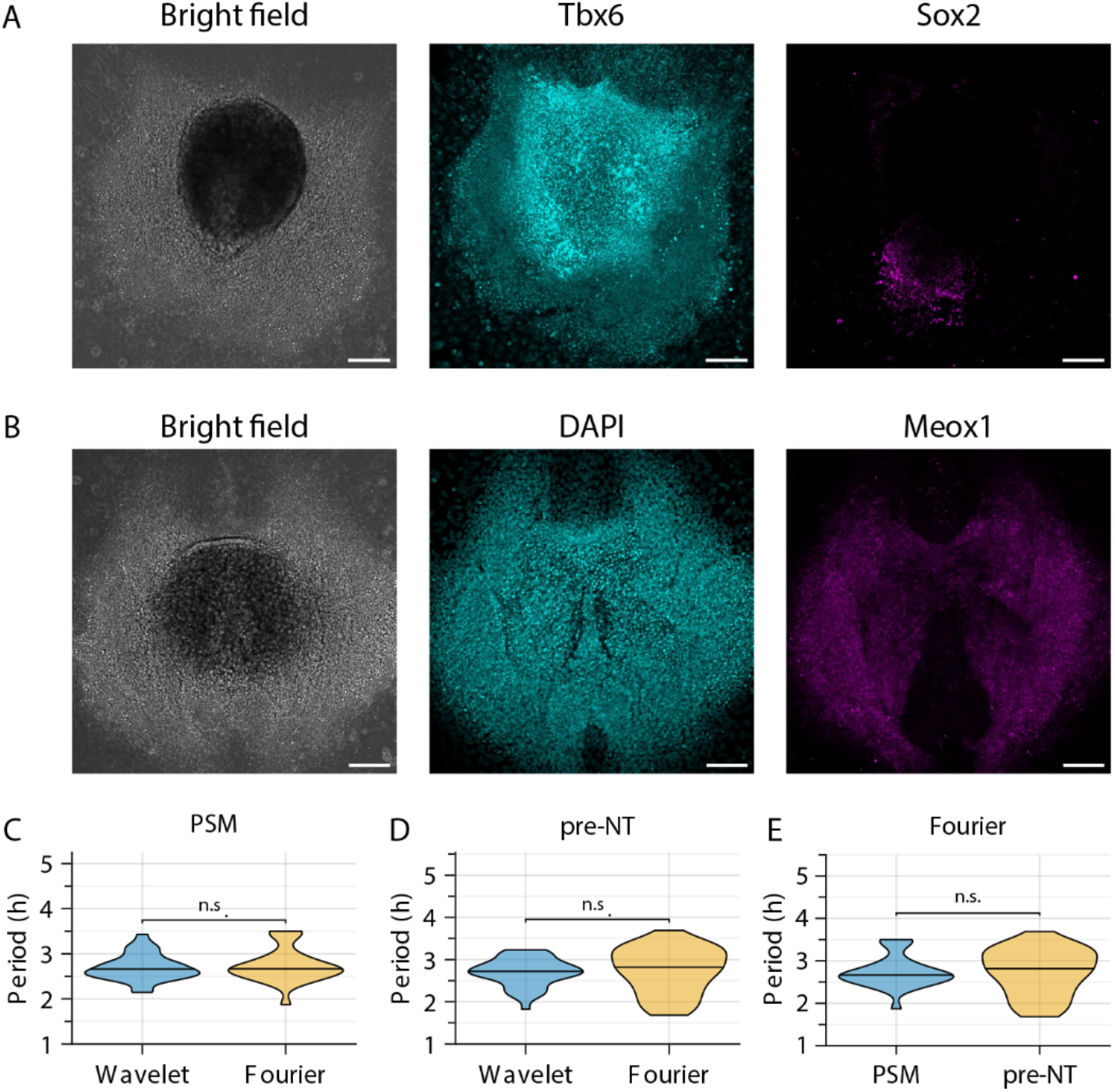
Characterization of 2D *ex vivo* cultures of E10.5 embryonic tail tips. **A, B** Cultures were fixed after 20 h and stained. **A** Representative image of immunostaining for PSM marker Tbx6 and neural marker Sox2. **B** Representative image of HCR for Meox1 and counterstaining for nuclei (DAPI). Scale bar is 100 µm. **C, D** Periods were quantified using wavelet transform or Fourier transform for PSM cells (**C**) or pre-NT cells (**D**). In **E** periods were quantified using Fourier transform for PSM cells and pre-NT cells. The coefficient of variation is 0.1 for PSM cells and 0.2 for pre-NT cells. * is p<0.05, ** is p<0.01, *** is p<0.001, **** is p<0.0001. PSM cells: n=39; pre-NT cells: n=40.

**Figure S5.**
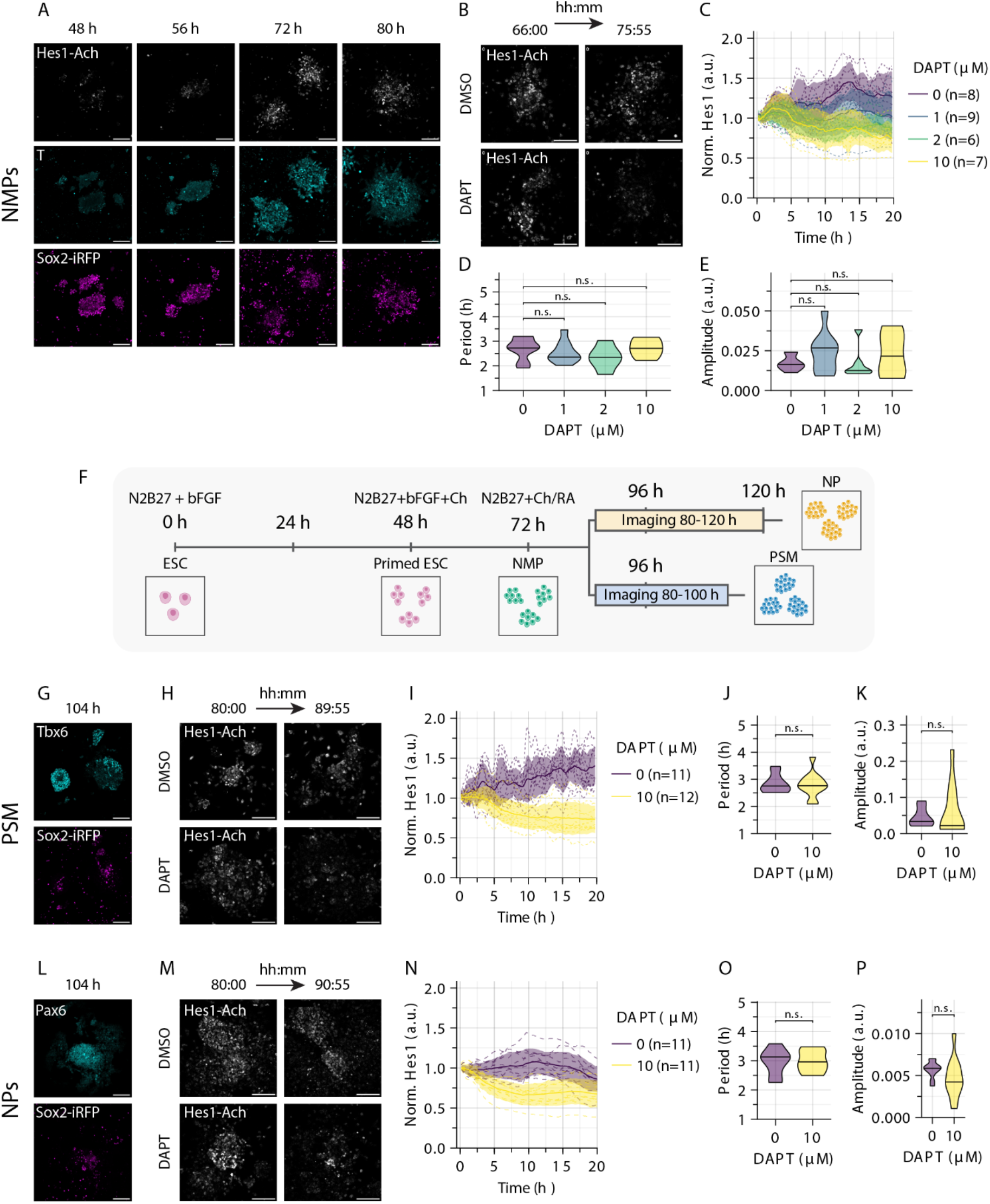
Population-wide Hes1 dynamics in *in vitro* differentiated cells. **A** Mouse ESCs expressing Hes1-Achilles and Sox2-iRFP were differentiated *in vitro* to NMPs. Cells were fixed at the indicated timepoints and immunostaining against T was performed. Representative images are shown. **B-E** Quantification of Hes1-Achilles dynamics in whole colonies. Note that data for control corresponds to data in Fig. 5. Representative snapshots are shown in **B**. *In vitro* differentiated NMPs were incubated with DMSO control or the gamma-secretase inhibitor DAPT. Hes1-Achilles intensity was measured over time. In **C** individual tracks (dotted lines), mean (solid line) and standard deviation (area) of the timeseries data (DMSO-normalised) are shown. Note the dose-dependent decrease in Hes1-Achilles intensity. Quantification of the period (**D**) and amplitude (**E**) by wavelet transform. **F-K** Hes1-Achilles dynamics in *in vitro* differentiated PSM colonies. **F** Mouse ESCs expressing Hes1-Achilles and Sox2-iRFP were differentiated *in vitro* along NMPs to PSM or neural progenitor (NP) colonies. Hes1-Achilles intensity was measured by fluorescence real-time imaging from 80 - 100 h of differentiation for PSM colonies and 80 – 120 h of differentiation for NP colonies. **G** Colonies were fixed at the indicated timepoints and immunostaining against Tbx6 was performed. Representative images are shown. **H-K** Quantification of Hes1-Achilles dynamics in PSM colonies. Representative snapshots are shown in **H**. *In vitro* differentiated PSM colonies were incubated with DMSO control or the gamma-secretase inhibitor DAPT. Hes1-Achilles intensity was measured over time. In **I** individual tracks (dotted lines), mean (solid line) and standard deviation (area) of the timeseries data (DMSO-normalised) are shown. Quantification of the period (**J**) and amplitude (**K**) by wavelet transform. **L** Cells were fixed at the indicated timepoints and immunostaining against Pax6 was performed. Representative images are shown. **M-P** Quantification of Hes1-Achilles dynamics in NP colonies. Representative snapshots are shown in **M**. *In vitro* differentiated NP colonies were incubated with DMSO control or the gamma-secretase inhibitor DAPT. Hes1-Achilles intensity was measured over time. In **N** individual tracks (dotted lines), mean (solid line) and standard deviation (area) of the timeseries data (DMSO-normalised) are shown. Quantification of the period (**O**) and amplitude (**P**) by wavelet transform. Scale bar is 100 µm.

**Figure S6.**
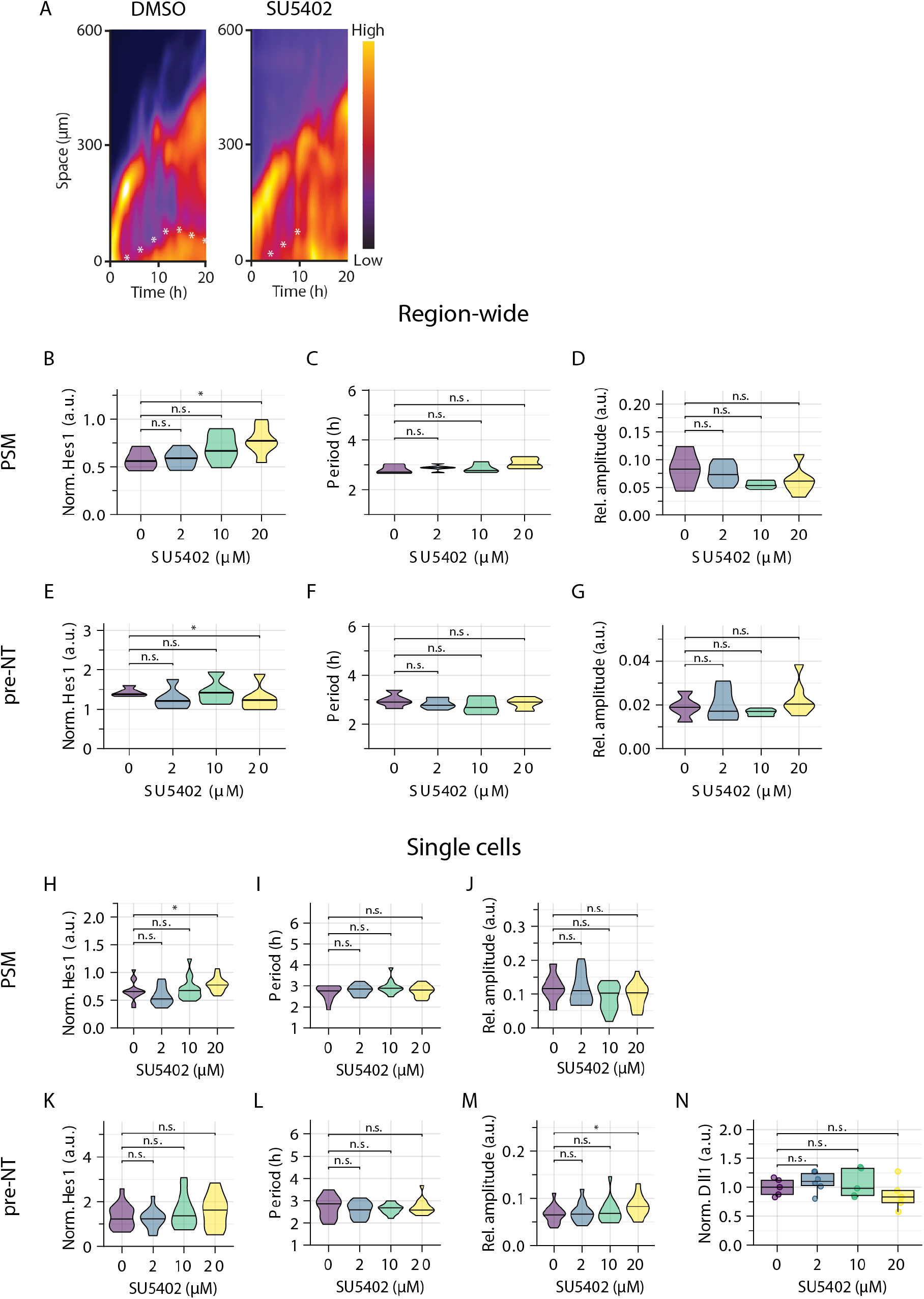
FGF inhibition leads to an increase in Hes1 oscillation amplitude in pre-NT cells. Further analysis of data shown in Fig. 6. **A** Representative kymographs for PSM region upon DMSO and FGF inhibition (SU5402) are shown. Hes1-Achilles wave is indicated by asterisk. **B-D** Quantification of population-wide dynamics in the PSM region (0 µM: n=5; 2 µM: n=6; 10 µM: n=5; 20 µM: n=7). **B** Quantification of absolute (DMSO normalised) Hes1-Achilles expression levels. **C, D** Quantification of period (**C**) and relative (mean-normalized) amplitude (**D**) by wavelet transform. **E-G** Quantification of Hes1-Achilles dynamics in pre-NT region (0 µM: n=5; 2 µM: n=6; 10 µM: n=5; 20 µM: n=7). **E** Quantification of absolute (DMSO normalised) Hes1-Achilles expression levels. **F, G** Quantification of period (**F**) and relative (mean-normalized) amplitude (**G**) by wavelet transform. **H-J** Quantification of Hes1-Achilles dynamics in PSM cells (0-10 µM: n=15; 20 µM: n=18). Quantification of absolute (DMSO normalised) Hes1-Achilles expression levels (**H**) and period (**I**) and relative (mean-normalised) amplitude (**J**) by wavelet transform. **K-M** Quantification of Hes1-Achilles dynamics in pre-NT cells (each concentration n=15). Quantification of absolute (DMSO-normalised) Hes1-Achilles expression levels (**K**) and period (**L**) and relative (mean-normalised) amplitude (**M**) by wavelet transform. **N** Cultures were fixed after 18 h and immunostaining against Dll1 and Sox2 was performed. Quantification of Dll1 in the Sox2-positive region is depicted (per concentration minimal n=5). Dots in boxplots represent individual data points. * is p<0.05, ** is p<0.01, *** is p<0.001, **** is p<0.0001.

**Figure S7.**
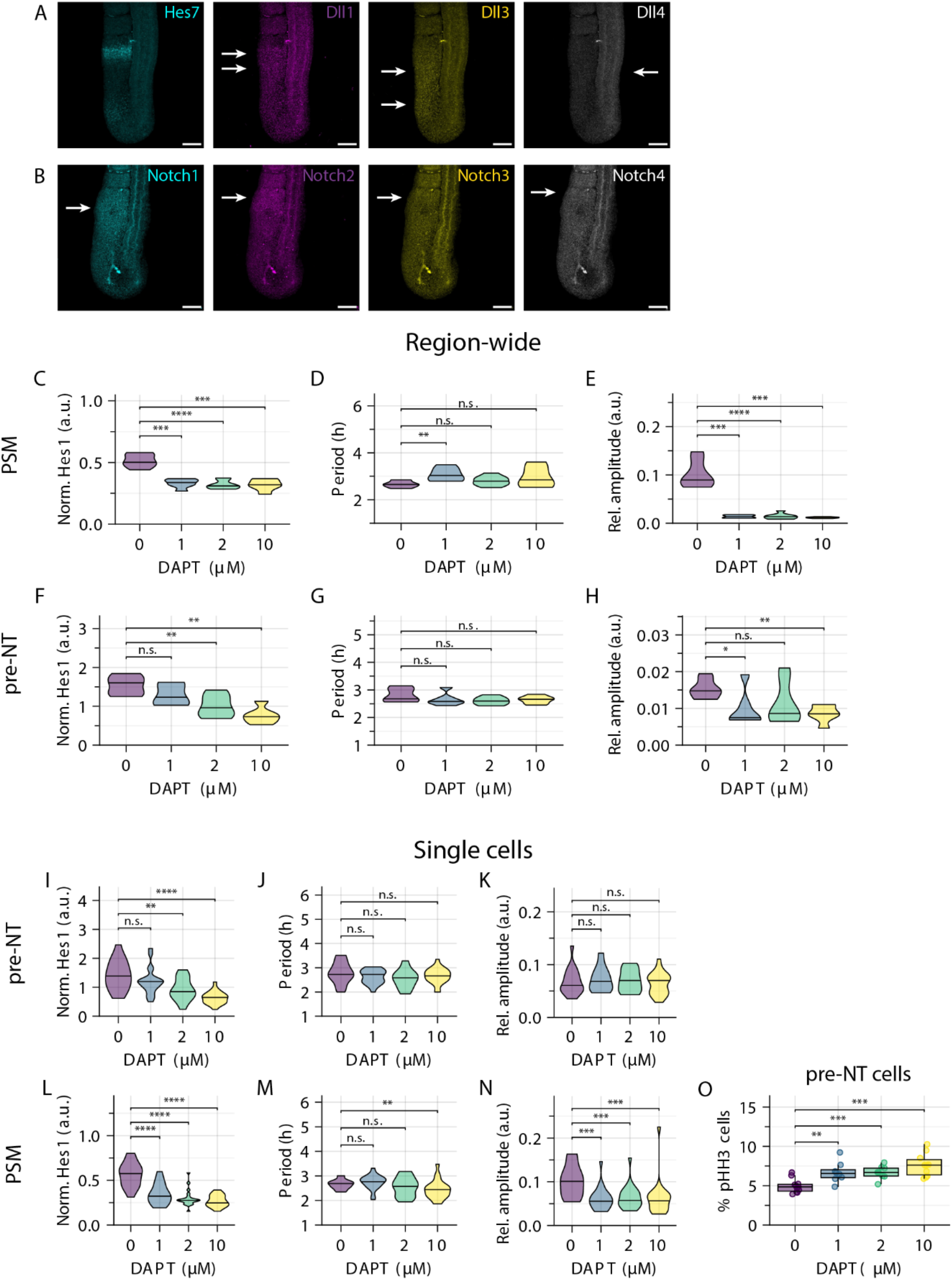
Notch inhibition has different effects on Hes1 dynamics in the pre-NT and PSM. **A, B** Expression of Delta-like ligands in comparison to Hes7 (**A**) and expression of Notch receptors (**B**) was visualized in E10.5 embryonic tails by HCR. Arrows highlight the expression pattern in the posterior embryonic tail. **C-O** Further quantification of the data shown in Fig. 7. E10.5 embryonic tails were cultured *ex vivo* as 2D cultures and treated with the indicated concentrations of the Notch inhibitor DAPT. **C-E** Quantification of population-wide dynamics in the PSM region (0 µM: n=8; 2 µM: n=6; 10 µM: n=8; 20 µM: n=6). **C** Quantification of absolute (DMSO normalised) Hes1-Achilles expression levels. **D, E** Quantification of period (**D**) and relative (mean-normalized) amplitude (**E**) by wavelet transform. **F-H** Quantification of population-wide dynamics in the pre-NT region (0 µM: n=8; 2 µM: n=6; 10 µM: n=8; 20 µM: n=6). **F** Quantification of absolute (DMSO normalised) Hes1-Achilles expression levels. **G, H** Quantification of period (**G**) and relative (mean-normalized) amplitude (**H**) by wavelet transform. **I-K** Quantification of Hes1-Achilles dynamics in single pre-NT cells (0 µM: n=18; 1 µM: n=16; 2 µM: n=21; 10 µM: n=21). Quantification of absolute (DMSO-normalised) Hes1-Achilles expression levels (**I**) and period (**J**) and relative (mean-normalised) amplitude (**K**) by wavelet transform. **L-N** Quantification of Hes1-Achilles dynamics in single PSM cells (0 µM: n=18; 1 µM: n=21; 2 µM: n=21; 10 µM: n=21). Quantification of absolute (DMSO-normalised) Hes1-Achilles expression levels (**L**) and period (**M**) and relative (mean-normalised) amplitude (**N**) by wavelet transform. **O** Cultures were fixed after 18 h and immunostaining against phosphorylated histone H3 (pHH3) and Sox2 was performed. Quantification of pHH3-positive cells in the Sox2-positive region is depicted (per concentration minimal n=8). Dots in boxplots represent individual data points. * is p<0.05, ** is p<0.01, *** is p<0.001, **** is p<0.0001.

