## Extended-data for "Unravelling differential Hes1 dynamics during axis elongation of mouse embryos through single-cell tracking"

**Data Figure 3. Timeseries data in regions of embryo tail cultures**

Normalized and detrended timeseries data of tails analyzed in Figure 3
blue: PSM, green: tailbud, yellow: pre-NT

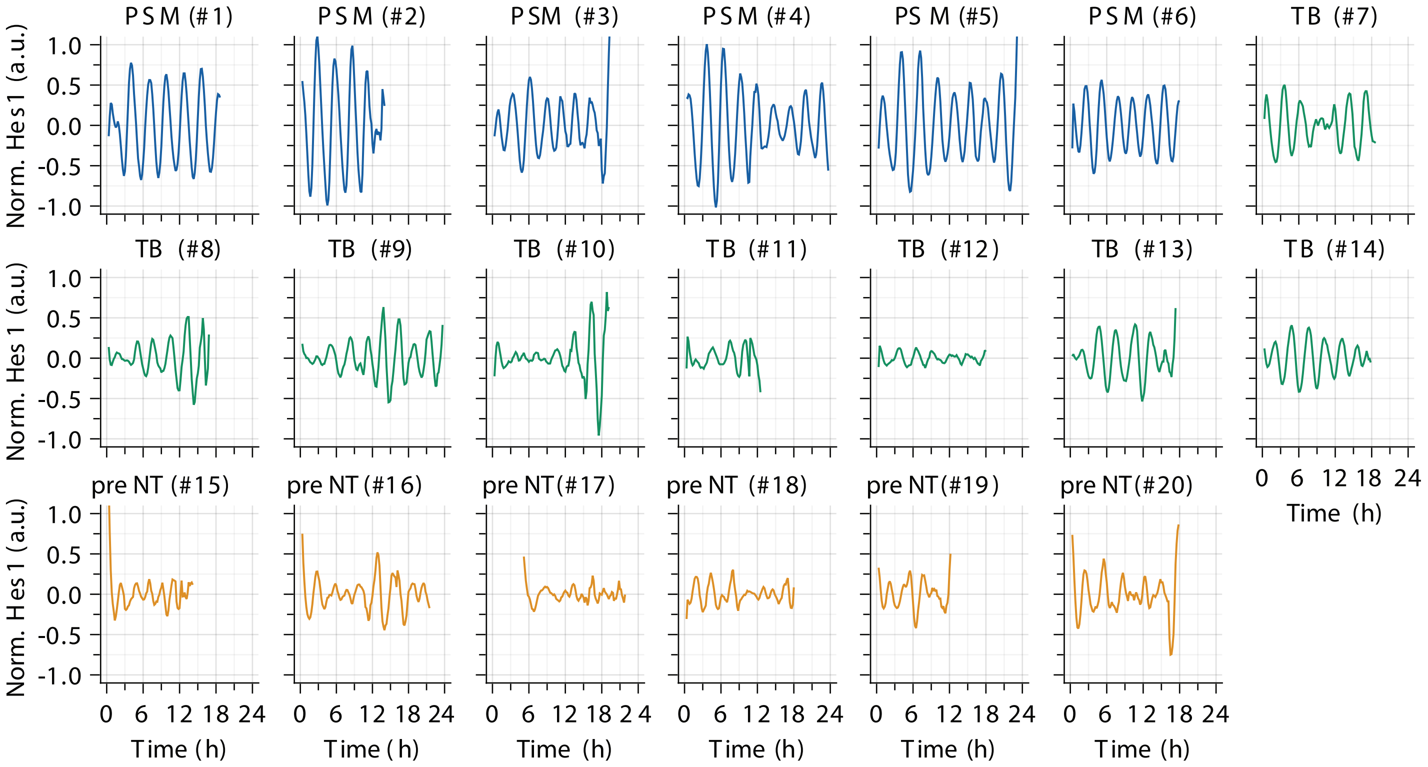

### Data Figure 4. Single cell tracks in spreadout cultures

Normalized and detrended timeseries data of single cells analyzed in Figure 4
blue: PSM, yellow: pre-NT

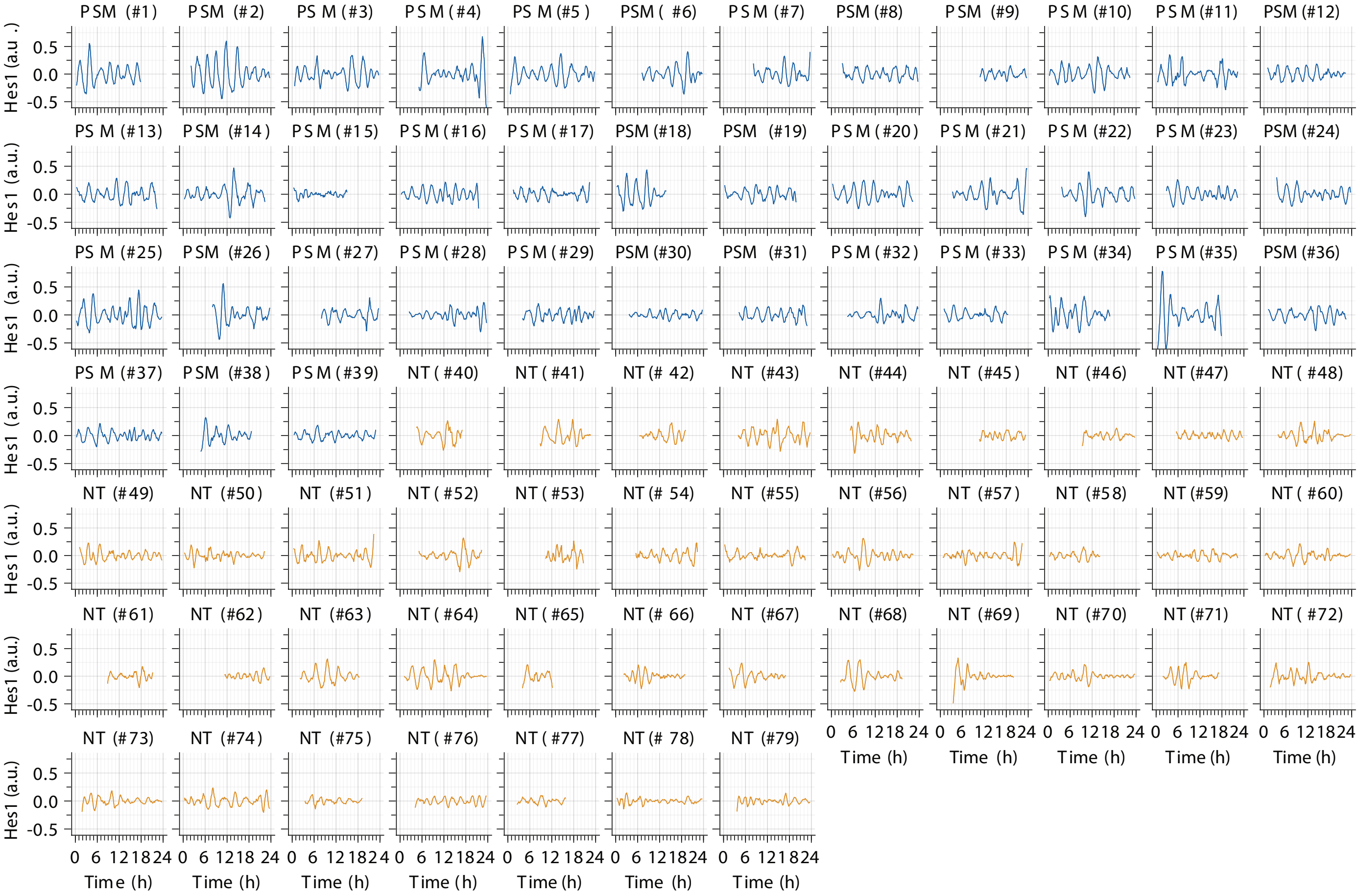


Fourier spectra of timeseries data of single cells analyzed in Figure 4
blue: PSM, yellow: pre-NT

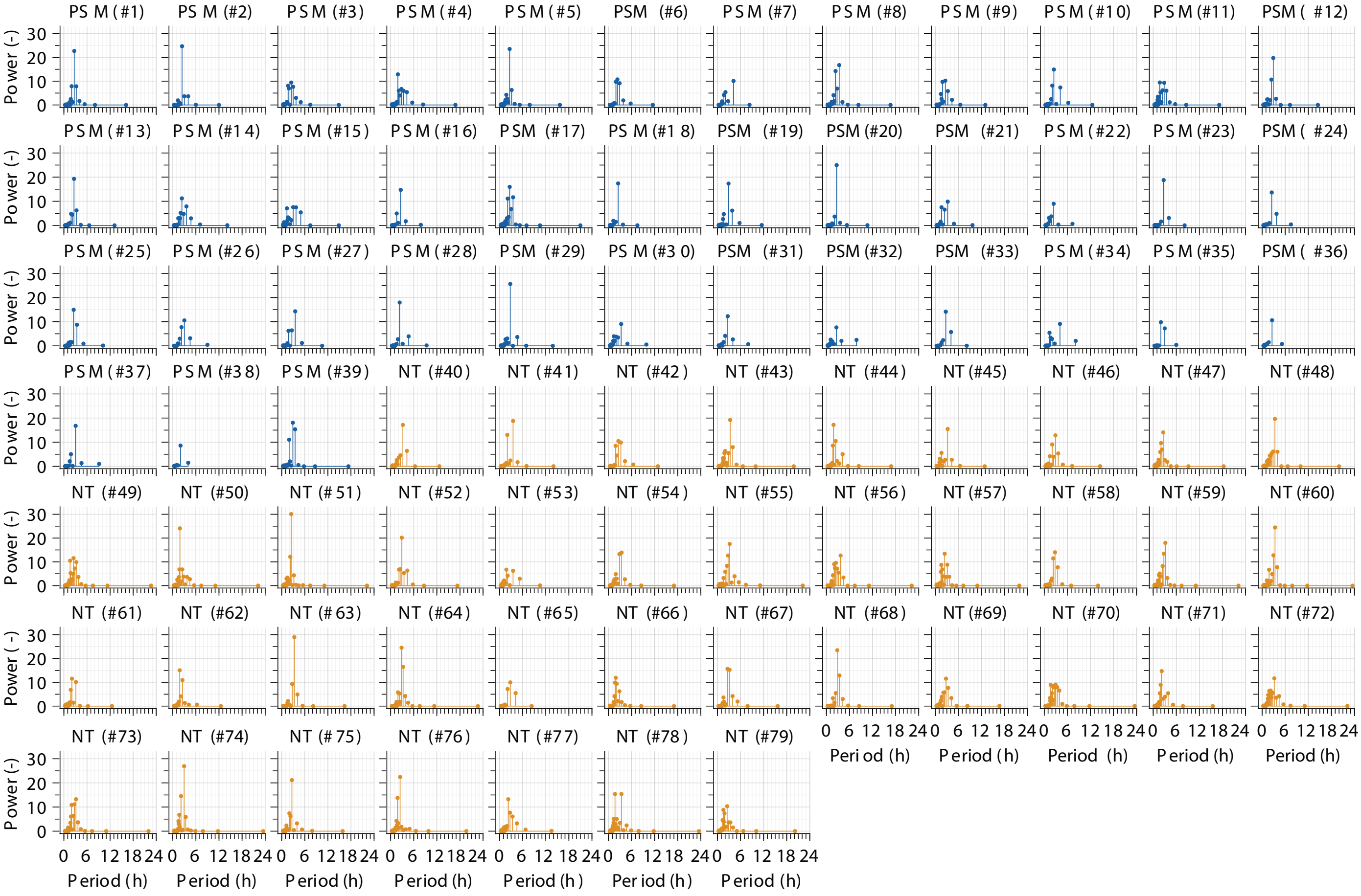

### Data Figure 6. Single cell tracks in spreadout cultures upon FGF inhibition

#### Normalized and detrended timeseries data of single pre-NT cells analyzed in Figure 6

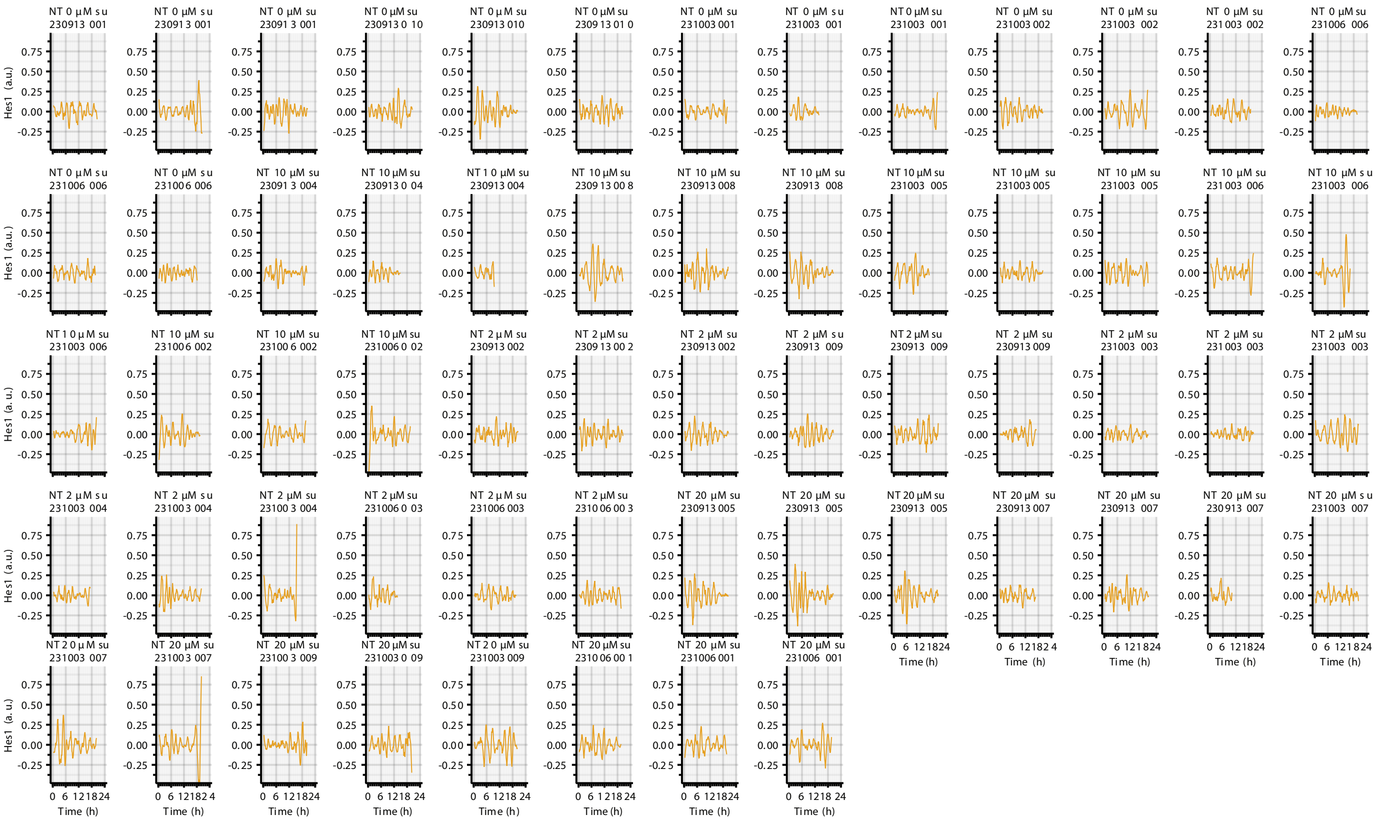


#### Normalized and detrended timeseries data of single PSM cells analyzed in Figure S6

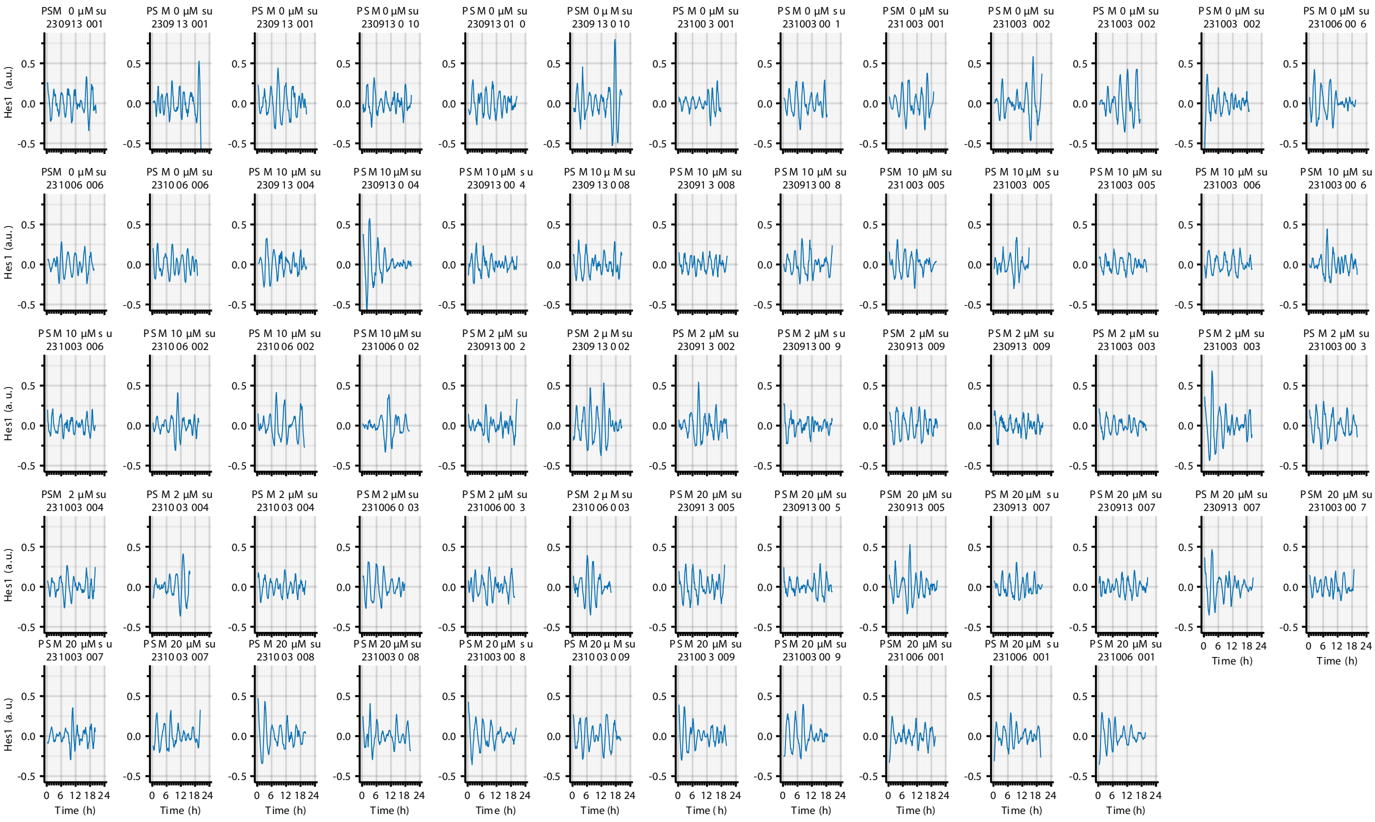

**Data Figure 7. Single cell tracks in spreadout cultures upon Notch inhibition**

#### Normalized and detrended timeseries data of single pre-NT cells analyzed in Figure 7

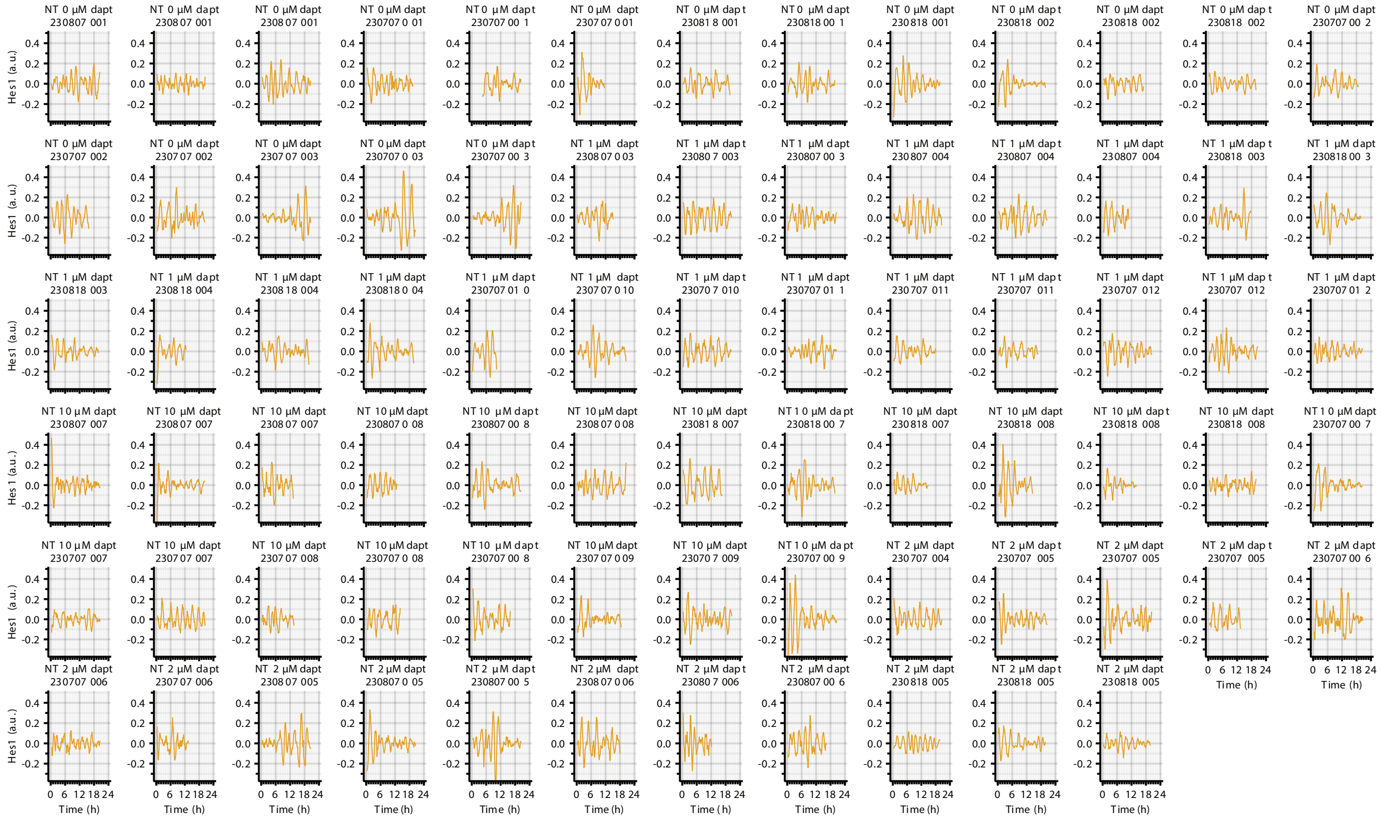

#### Normalized and detrended timeseries data of single PSM cells analyzed in Figure S7

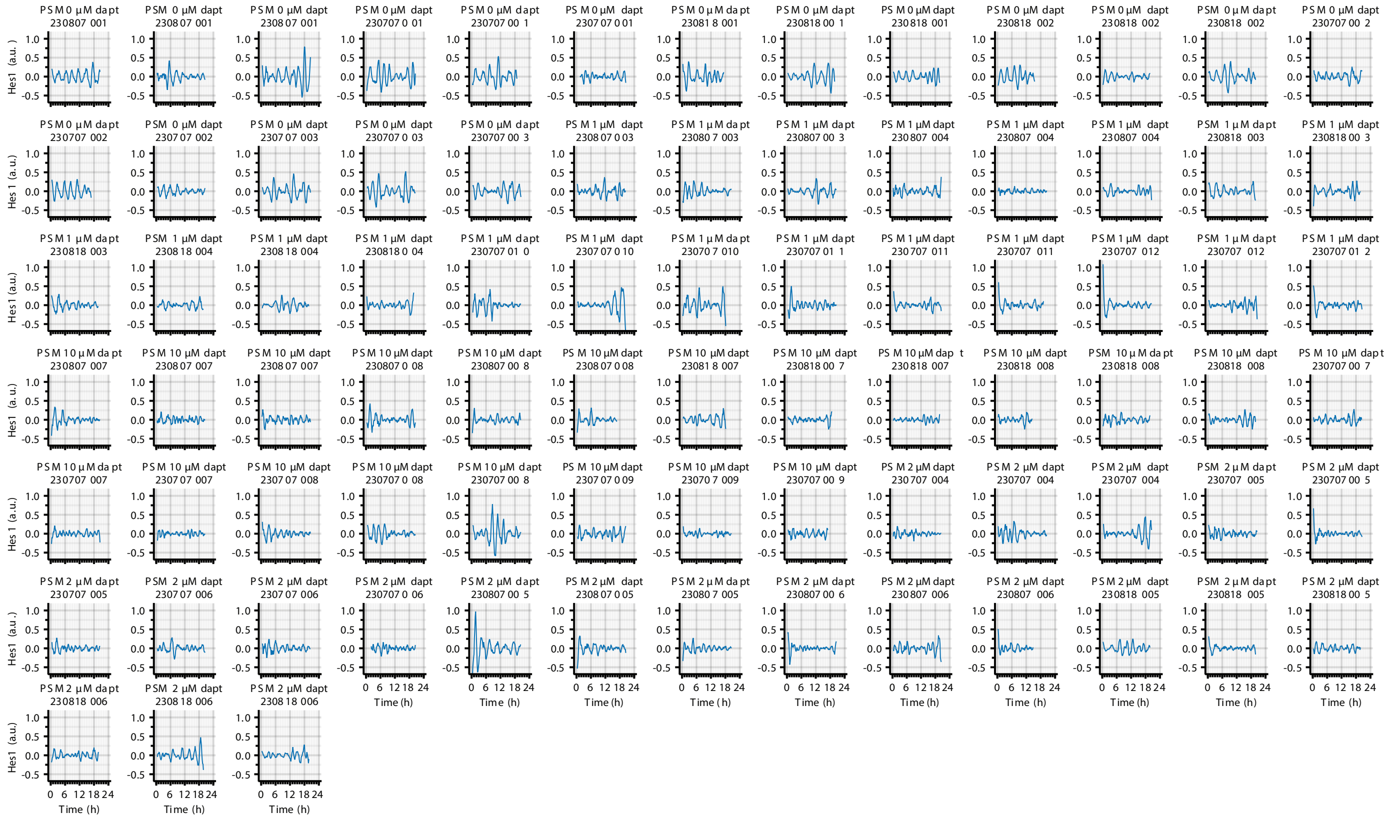
